# AF-vapeR: A multivariate genome scan for detecting parallel evolution using allele frequency change vectors

**DOI:** 10.1101/2021.09.17.460770

**Authors:** James R. Whiting, Josephine R. Paris, Mijke J. van der Zee, Bonnie A. Fraser

## Abstract

1. The repeatability of evolution at the genetic level has been demonstrated to vary along a continuum from complete parallelism to divergence. In order to better understand why this continuum exists within and among systems, hypotheses must be tested using high confidence sets of candidate loci for repeatability. Despite this, few methods have been developed to scan SNP data for signatures specifically associated with repeatability, as opposed to local adaptation.
2. Here we present AF-vapeR (Allele Frequency Vector Analysis of Parallel Evolutionary Responses), an approach designed to identify genome regions exhibiting highly correlated allele frequency changes within haplotypes and among replicated allele frequency change vectors. The method divides the genome into windows of an equivalent number of SNPs, and within each window performs eigen decomposition over normalised allele frequency change vectors (AFV), each derived from a replicated pair of populations/species. Properties of the resulting eigenvalue distribution can be used to compare regions of the genome for those exhibiting strong parallelism, and can also be compared against a null distribution derived from randomly permuted AFV. Further, the shape of the eigenvalue distribution can reveal multiple axes of parallelism within datasets.
3. We demonstrate the utility of this approach to detect different modes of parallel evolution using simulations, and also demonstrate a reduction in error rate compared with intersecting F_ST_ outliers. Lastly, we apply AF-vapeR to three previously published datasets (stickleback, guppies, and Galapagos finches) which comprise a range of sampling and sequencing strategies, and lineage ages. We detect known parallel regions whilst also identifying novel candidates.
4. The main benefits of this approach include a reduced false-negative rate under many conditions, an emphasis on signals associated specifically with repeatable evolution as opposed to local adaptation, and an opportunity to identify different modes of parallel evolution at the first instance.

## INTRODUCTION

Studies of convergence, where separate lineages evolve similar phenotypes, provide compelling evidence of natural selection and have been a prominent focus of research in the last decade (Stern 2013; Storz 2016; Blount, Lenski, and Losos 2018; Arendt and Reznick 2008). Common practice in population genomics studies of repeatability involves sequencing populations in replicated environments, performing selection scans, and comparing candidate loci across replicates (the overlapping outlier approach reviewed in (Fraser and Whiting 2019)). Identified selection candidates are then probed for theories of genetic convergence; for example their mode of convergence (Lee and Coop 2017), the role of genetic and pleiotropic constraints (Morris et al. 2019; Wollenberg Valero 2020), the influence of redundancy in the genotype-phenotype map (Yeaman et al. 2018; Láruson, Yeaman, and Lotterhos 2020), or the effect of gene flow (Fang et al. 2021). It is also of interest whether repeatability occurs through: 1) selection at the same SNP (often termed ‘parallel’ evolution), as seen in stickleback (Colosimo 2005) and maize (L. Wang et al. 2021); 2) nonidentical changes in the same genes or loci (‘convergence’) e.g. repeated modification of *Mc1r* during pigmentation evolution (Manceau et al. 2010); 3) changes in different genes of similar function (‘functional convergence’) e.g. developmental HOX genes across aquatic mammals (Nery et al. 2016); or 4) using no comparable genetic machinery whatsoever (‘divergence’).

Genome scans are a fundamental tool in identifying locally-adapted genomic regions, but scrutiny has increased in response to concerns of false-positives, biases, and interpretation of outliers (Booker et al., 2021; Cruickshank & Hahn, 2014; Lotterhos & Whitlock, 2015). False-positives may be non-randomly associated with shared recombination landscapes between lineages (Booker et al., 2020; Lotterhos, 2019), but random false-positives could be due to drift (Leigh et al., 2021), or a tendency towards detecting hard-sweeps over soft-sweeps (Teshima et al., 2006). Generally, the overlapping outlier test assumes false-positives are randomly distributed and they should not overlap among replicate lineages. However, over-conservative cut-offs or structure corrections subsequently may mask overlapping true positives and inflate the false-negative rate (FNR) in convergence studies. This implies that some mechanisms of repeatable adaptation may be more difficult to detect, which is concerning given most adaptation may occur through minor/moderate allele frequency change (soft sweeps) (Hermisson & Pennings, 2017).

Many definitions of convergence/parallelism exist (Stuart, 2019), including those which consider these as geometric processes in multivariate space (Adams & Collyer, 2009; De Lisle & Bolnick, 2020). Applying these here, ‘convergence’ is a process where distinct evolutionary starting points converge on a common endpoint. ‘Parallelism’ is a process where distinct evolutionary start points change in the same way along parallel trajectories to arrive at distinct endpoints. Parallel lineages, therefore, may not evolve to become more similar, contrary to convergent lineages. At the genetic level, parallelism can be reflected by changes in allele frequencies in the same direction at the same sites. We can then also define ‘anti-parallelism’ as changes in allele frequencies in the opposite direction at the same sites, and ‘non-parallelism’ as changes in allele frequencies at different sites within the same genomic regions. ‘Multi-parallelism’ may also exist, where multiple parallel trajectories, each with ‘parallel’ lineages, are observed in the same genomic region. Multi-parallelism can therefore be analogous to definitions of ‘convergence’ involving evolution at different sites affecting the same gene or locus. Multi-parallelism therefore represents special instances of non-parallelism, in which several non-parallel axes exist, each with multiple lineages evolving in parallel on a given axis. These various forms of parallelism in allele frequency change (summarised in Figure S1) will allow us to identify parallelism, anti-parallelism, and multi-parallelism, within genomes.

We present a novel population genomics scan to specifically identify regions associated with genetic parallelism, bypassing the requirement to first identify local adaptation candidates in replicate datasets. We first outline the background of this approach in detail, before demonstrating its usage and comparing it to F_ST_-based overlapping outlier approaches using simulations, and finally apply it to a range of empirical datasets of different sampling design, sequencing resolution, and biological questions. We have released this method as an R package (https://github.com/JimWhiting91/afvaper), which we expect to become an important tool for researchers interested in identifying genomic regions associated with genetic parallelism.

## MATERIALS AND METHODS

### Allele Frequency Vector Analysis of Parallel Evolutionary Responses (AF-vapeR)

Our method is heavily inspired by the work of De Lisle and Bolnick (2020), and extends their framework for deriving a single phenotypic eigenvalue distribution to many allele frequency-derived eigenvalue distributions across the genome. These are then compared against randomly permuted distributions across the genome.

SNP data is first split into non-overlapping windows (Figure 1A) of equal user-defined SNP count, within which we calculate per-SNP allele frequency change between multiple population pairs. Population pairs are defined based on *a-priori* knowledge of the system, such that each pair represents a vector of change from ecotype/phenotype/habitat A to ecotype/phenotype/habitat B. These could be unique ‘replicate pairs’ (each A-B population is unique); a single A to many B (‘common’ ancestor); or vectors between a pool of A and B populations sampled with replacement (‘multi-comparisons’). These setups reflect sampling designs common to studies of genetic convergence (Fraser & Whiting, 2019).

**Figure 1:**
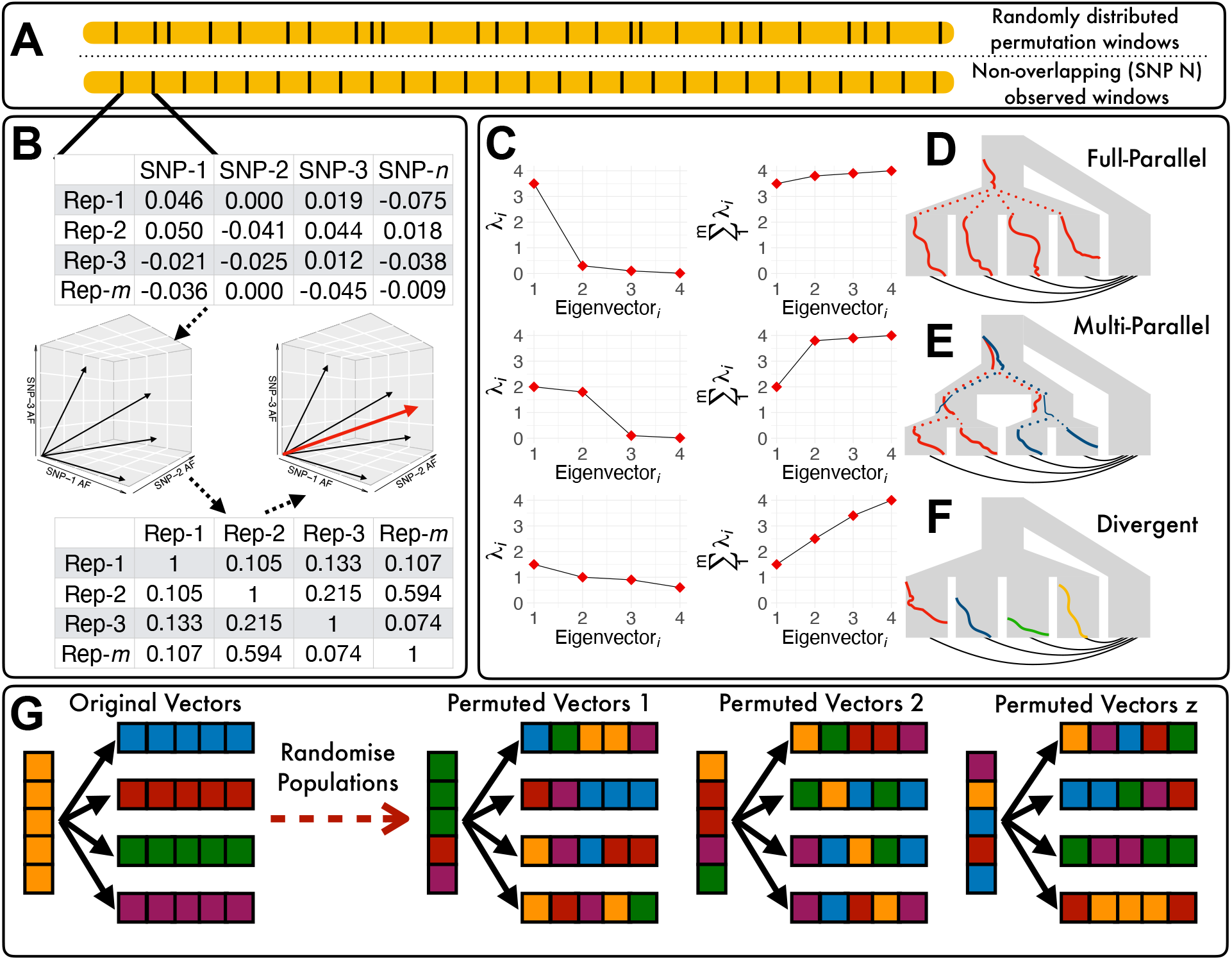
Methodology of AF-vapeR. Chromosomes are windowed by SNP count (**A**). Normalised AFV are calculated for all population pairs within a window (**B**) prior to calculating the correlation matrix and eigen decomposition to yield eigenvectors (red arrow as example). Eigenvalues (λ) are summed along consecutive eigenvectors (1.m) (**C**), with large summed values highlighting that a majority of variance is constrained to a few eigenvectors; indicative of parallelism. Distribution shape is indicative of different evolutionary processes (**D-F**), with large primary eigenvectors highlighting full parallelism, large second eigenvalues highlighting two axes of parallelism, and no large eigenvalues suggesting divergence/non-parallelism. Different evolutionary processes are shown as increasing in frequency from left to right within lineages. Dotted lines show inheritance through lineage branching. Colours denote shared mutations, and black lines connecting lineages show AFV pairs. The permutation process is illustrated at the bottom (**G**), where fixed population assignment, denoted as block colour, is repeatedly shuffled and AFV are calculated under the same vector design for a total of z permutations. Individuals are squares and each column/row represents a population, with AFV shown as black arrows between the vertical population to all four horizontal populations in a common-ancestor design.

The process produces a series of *m* x *n* matrices, where *m* represents the number of population pairs and *n* represents the number of SNPs within each window (Figure 1B). Each cell therefore describes the change in allele frequency (ΔAF) for a single SNP from population *P_a_* to *P_b_* (equation 1). The vector *P_a_* → *P_b_* describes ΔAF for all SNPs in the window for a given population pair; termed the allele frequencies vector (**AFV**) (equation 2). The final matrix **X** includes the normalised **AFV** for all population pairs 1.*m*, where normalisation involves dividing each vector by its length to a unit length of one (equation 3), such that all population pairs exhibit equivalent variance. As such, the absolute extent of allele frequency change is only relevant between SNPs within a window, within a population pair (i.e. within each **AFV**).

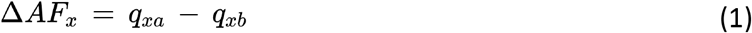

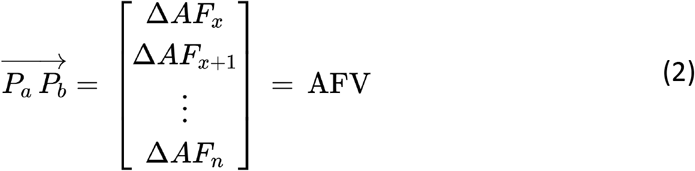

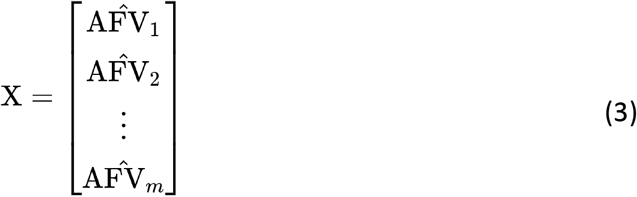

For each **X** matrix, we then calculate matrix **C,** which is an *m* x *m* matrix describing the correlations between normalised **AFV** within **X** (equation 4), and subsequently perform eigen decomposition of **C** to examine parallelism within the focal genomic window (equation 5). Here, **Q** is the matrix of eigenvectors and **V** is the diagonal matrix of eigenvalues, as described in De Lisle and Bolnick (2020).

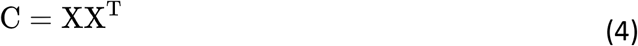

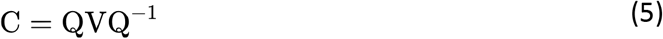

Because the **AFV** are normalised to unit length, all eigenvalues in a window sum to *m*, so we are interested in identifying regions of the genome in which the eigenvalue distribution is positively skewed and weighted to variance captured by only the first (or first few) eigenvalues (Figure 1C). The raw eigenvalues associated with the *m*-1 eigenvectors can thus be compared and visualised across the genome. The eigenvectors of **C** can also be related back to the allele frequency change at individual SNPs according to equation 6.

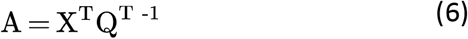

To develop a null expectation for extreme eigenvalues, we repeatedly reassign individuals to random populations, maintaining the original vector design strategy (Figure 1G), and recalculate random **AFV** across the genome. This approach captures potential biasing of eigenvalues across regions of the genome, for example due to recombination and linkage variation, and bias associated with vector sampling design that would otherwise fail to be accounted for by a one-size-fits-all null model such as the Wishart distribution (De Lisle & Bolnick, 2020).

### Simulation methods

Simulation methods are described in full in the supplementary materials. Briefly, to demonstrate functionality and explore the effectiveness of AF-vapeR over parameter space, we used a Wright-Fisher model with the forward-simulation software SLiM (v3.4 (Haller & Messer, 2019)). The goals of these simulations were to: 1) Demonstrate the power to detect parallelism under different demographies and selection contexts; 2) Demonstrate the ability to detect different evolutionary scenarios (Figure 1D-F); and 3) Compare *AF-vapeR* to the commonly applied approach of overlapping F_ST_ outlier windows among replicates.

We simulated a five-population metapopulation with an ancestral founding population (N=10,000) and four daughters (N=400, 250, or 10,000). After founding, each daughter experienced 200 generations of selection on an ancestrally-neutral mutation inherited at low-frequency from the ancestral population. Either: 1) all daughters inherited and swept the same mutation (full-parallelism); 2) daughters were paired, and each pair swept their pair-specific mutations (multi-parallelism); or 3) all daughters swept their own mutation (non-parallelism/divergence) (Figure S2). We manipulated the metapopulation by modifying: the size of all daughter populations (to encourage drift), the reciprocal migration rate between the ancestral and all four daughters (*M* = 0, 0.0025, 0.01), and the selection coefficient applied to the focal mutation in all daughters (*s* = 0.01, 0.05, 0.1). Factorially combining these parameters yielded 27 treatment groups. Separately, we explored how positioning the focal mutation in regions of low, medium, and high recombination, affected results.

AF-vapeR (window sizes of 50, 200, 500 SNPs) was compared to F_ST_ scans (10kb windows) in false-positive rate (FPR) and false-negative rate (FNR) with variable cutoffs (α = 0.95, 0.99, 0.999). FPR reflected the proportion of falsely-identified windows beyond the selected mutation averaged across all 100 iterations. FNR reflected the number of iterations that failed to identify parallelism at the focal mutation. All simulation parameters are in Table S1.

### Empirical dataset demonstrations

We chose three publicly available datasets to demonstrate different utilities of AF-vapeR. These include restriction-site associated digestion (RAD) sequencing data from 18 freshwater-adapted three-spined stickleback populations British Columbia (BC) and an ancestral marine population (Magalhaes et al., 2016, 2020); whole-genome sequencing (WGS) of five pairs of high-predation (HP) and low-predation (LP) adapted guppies (Whiting et al., 2021); and WGS data from 12 species of Galapagos finches, used previously to explore small vs large, and pointed vs blunt, beak morphologies within the adaptive radiation (Han et al., 2017; Lamichhaney et al., 2015, 2016). These datasets therefore include different sequencing approaches (RAD-seq vs WGS), different sampling approaches (replicate pairs vs common ancestor vs multiple comparisons; see (Fraser & Whiting, 2019)), and a mix of within- and across-species comparisons.

## RESULTS

### Simulation results

AF-vapeR was successfully able to distinguish between the three different simulation scenarios: Full Parallelism (Figure 2A), Multi-Parallelism (Figure 2B), and Divergent (Figure 2C). These distinctions can be seen as clear peaks arising on the first (Full) or second (Multi) eigenvector, or not at all (Divergent) at the selected site at 10Mb.

**Figure 2:**
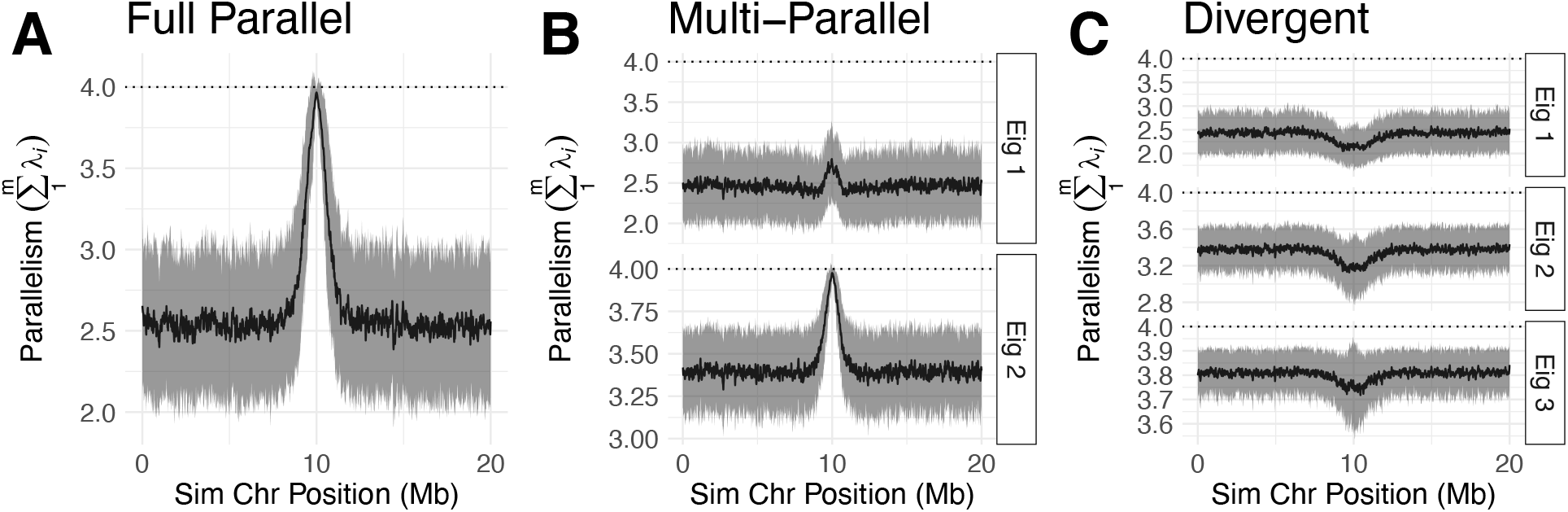
Simulated eigenvalue peaks (with standard deviation) detected by AF-vapeR under three different evolutionary scenarios: full parallelism (**A**), multi-parallelism (**B**), and divergence (**C**). The maximum eigenvalue is 4. For each scenario we show the eigenvalues up until a peak is observed. The simulation parameters shown are for s = 0.05, M = 0.025, and DP = 10,000.

We then looked at error rates associated with detecting full parallelism. Across parameter space, at α = 0.01, FPR were low for all AF-vapeR window sizes (50 SNPs, max FPR = 0.057; 200 SNPs, max FPR = 0.072; 500 SNPs, max FPR = 0.085) and all overlapping F_ST_ thresholds (all 4 daughters, max FPR = 1.1e-4; any 3, max FPR = 7.5e-4; any 2, max FPR = 4.1e-3) (Figure S3). Low F_ST_ FPR suggests false-positives are randomly distributed within daughters. AF-vapeR FPR was positively associated with daughter population sizes and negatively associated with migration rate (Figure S3).

Across parameter space, FNR for all AF-vapeR window analyses consistently out-performed the strictest and most applicable F_ST_ analysis (outlier in all 4 daughters), and often out-performed all F_ST_ overlap cut-offs (Figure 3). Expectedly, strong bottlenecks (*DP* = 400, first column of panels in Figure 3), and strong migration (*M* = 0.01, first row of panels in Figure 3) led to elevated FNR for F_ST_ analyses, but AF-vapeR was generally more robust to these parameters.

**Figure 3:**
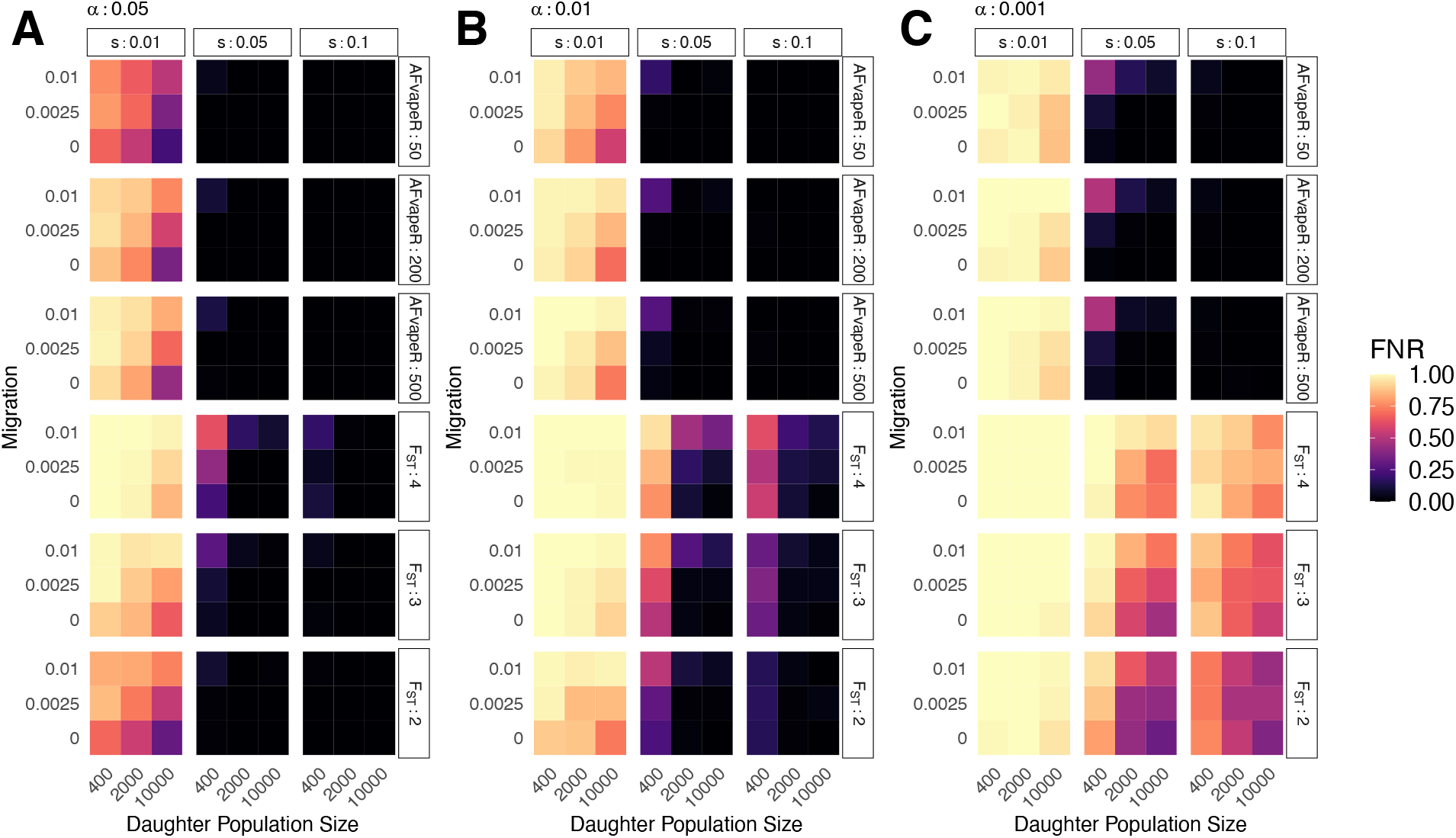
FNR associated with AF-vapeR and F_ST_ outliers tests. Facet labels correspond to AF-vapeR window size (50, 200, and 500 SNPs) or number of overlapping populations for F_ST_ (i.e. 2 = an outlier in any 2 of the 4 populations, 3 = an outlier in any 3, 4 = outlier in all four). Panels **A**, **B** and **C** show FNR calculated using quantile cutoffs of 95%, 99%, and 99.9% respectively. FNR was averaged over 100 iterations within cells. Facet columns = strength of selection on beneficial mutation, y = migration rate, and x = daughter population size.

For all F_ST_ overlapping cut-offs, weak selection (*s* = 0.01) often prevented identification of full-parallelism. Even using the least stringent cut-off, only 15% of simulations detected full-parallelism (α = 0.05, *s* = 0.01, F_ST_ outlier in all 4; min FNR = 0.85, max FNR = 1.0). In contrast, under the same conditions AF-vapeR recovered full parallelism in at least 21% and at most 78% of iterations under all nine parameter combinations of daughter population size, migration and *s* = 0.01 (α = 0.05, *s* = 0.01, AF-vapeR window size = 50; min FNR = 0.22, max FNR = 0.79). Detecting parallelism candidates as F_ST_ outliers in any two daughters improved FNR under weak selection (α = 0.05, *s* = 0.01, F_ST_ outlier in any 2; min FNR = 0.3, max FNR = 0.86). This approach, however, still does not perform as well as AF-vapeR and fails to capture the simulation conditions of parallel selection in all four daughters.

Under stronger selection (*s* = 0.05, 0.1), AF-vapeR exhibits consistently low FNR even as cut-off stringency increases (Figure 3A-C). Conversely, increasing cut-off stringency dramatically increases F_ST_-based FNR across all parameter space. This behaviour of AF-vapeR is important because it indicates that FPR at weak cut-offs can be ameliorated with stricter cut-offs at a minimal cost to false-negatives. Only with high migration and small population sizes (top left of parameter grids) did FNR increase with outlier stringency. Researchers should therefore consider the role of migration and population size when applying the method under such scenarios.

AF-vapeR was found to be more sensitive to lower allele frequency differentiation (AFD), as FNR reduced more rapidly with increasing AFD (Figure S4), which may explain the observed reduction in FNR when migration is high and selection is weak (low AFD). The reduced FNR seen when drift is strongest (DP = 400) is better explained by random drift at neutral sites reducing the background signal of parallelism and making peaks at selected sites more distinct.

Overall, these results show that AF-vapeR has comparably low FPR to overlapping multiple F_ST_ outlier sets, and has a generally reduced FNR across parameter space. AF-vapeR exhibits a considerably reduced FNR under drift, weak selection and high migration. AF-vapeR consistently outperformed F_ST_ when F_ST_ outliers must be observed in all four daughters, and thus performs better at capturing the simulation conditions of all daughters experiencing parallel selection. FNR were typically reduced with reduced window size for AF-vapeR with only small increases in FPR, suggesting the effect of noise is marginal when balancing signal and noise.

Comparable results were observed when running the simulation under a multi-parallelism scenario when looking for outliers at the selected site on the second eigenvector. In these simulations, as above, FPR was generally low (Figure S5), migration rate was positively associated with FNR, and selection coefficient was negatively associated with FNR (Figure S6). Cut-off stringency also behaved similarly, obscuring F_ST_-based analyses at higher stringencies. These results demonstrate that AF-vapeR is able to robustly identify parallelism candidates across parameter space whilst also inferring that multiple axes of parallelism exist.

Increasing recombination from 1e-9 to 1e-7 cM/bp increased the width of eigenvalue peaks and marginally reduced peak-height (Figure S7A). This was not affected by window SNP count. Eigenvalue variance at neutral sites increased by approximately 50% with a 100-fold reduction in recombination (Figure S7B-C), however peaks of full-parallelism were still clearly observable relative to the neutral backdrop regardless of local recombination rate.

### Empirical Results: Marine-Freshwater stickleback

The goal for this dataset was to detect *a-priori* loci associated with parallel evolution from RAD-sequencing data. We applied AF-vapeR to a dataset of 18 freshwater-adapted populations and a marine population in a ‘common-ancestor’ sampling strategy (Figure 4A) sampled from the northeast Pacific coast in British Columbia (Magalhaes et al., 2020) (Eighteen marine-freshwater AFV). Due to low SNP density (total = 26,658), we used windows of 10 SNPs (N=2376) with a median physical window size of 116.0 kb. At this resolution, the genomic landscape of full-parallelism, defined by the per-window primary eigenvalues (Figure 4B), closely resembled the genomic landscape of marine-freshwater divergence commonly observed in this system (Fang et al., 2021; Hohenlohe et al., 2010; Roberts Kingman et al., 2021). In particular, chromosome 4 exhibited a clear chromosome-level increase in parallelism (Figure 4C-D). Chromosomes 21, 12, 20 and 7 exhibited the next highest median primary eigenvalues (Figure 4D). With the exception of chr12, each of these are enriched for QTL associated with marine-freshwater phenotypes (Peichel & Marques, 2017).

**Figure 4:**
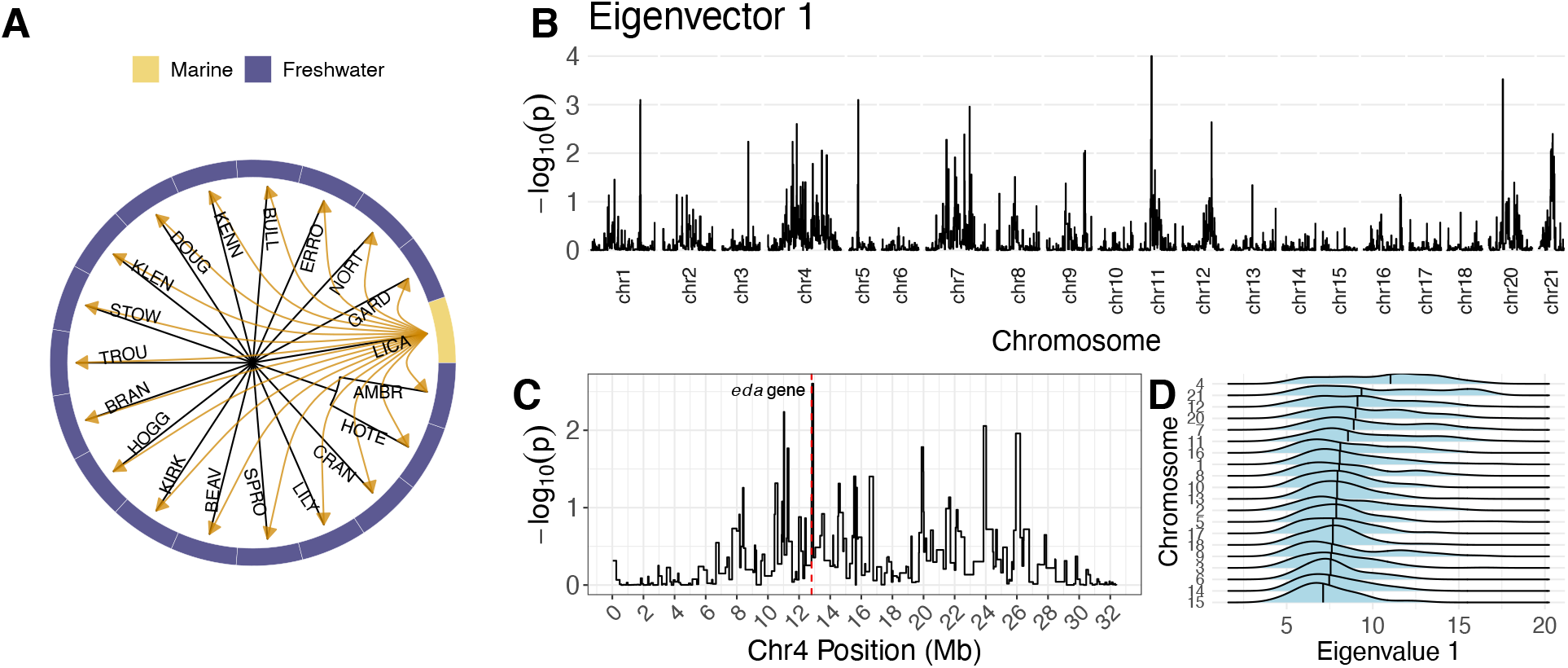
AF-vapeR results using a RAD-seq dataset for marine-freshwater stickleback from British Columbia. Sampling design is described based on the tree (**A**), with all vectors calculated from the marine population (LICA) to all freshwater populations. The genome-wide distribution of eigenvalue 1 is plotted in panel **B**, based on empirical p-values compared to ~10,000 null permutations. Chromosome 4 is highlighted in panel **C**, with the eda gene represented by a red dashed line. Per-chromosome distributions of eigenvalue 1 results are plotted as ridges in panel **D**, with the median denoted in each. Chromosomes are ordered vertically in terms of median strength of eigenvalue 1.

Above the permuted 99.9% cutoff, we detected six windows exhibiting full parallelism on chromosomes 1, 5, 7, 11, and 20, with the window on chromosome 11 exceeding all null permutations. All windows included previously identified candidates for freshwater adaptation (discussed in full in the supplementary results). Above the 99% cutoff, we recovered an additional 17 windows, including the window adjacent to the well-known *eda* gene associated with parallel marine-freshwater armour haplotypes. The window containing *eda* exhibited a weaker signal of full parallelism, likely because three of the 18 freshwater populations exhibit full-armour plating associated with marine fish (Magalhaes et al., 2020). Ten windows exceeded the 99% cutoff on eigenvector 2; none were above 99.9%. For these windows, eigenvalue 1 made up the majority of the summed total, with eigenvalue 2 explaining 15.32% or less of the total variance among AFV. This indicates that these windows represent cases in which most, but not all, freshwater populations are diverging in parallel on eigenvector 1, but additional axes exist to explain non-parallel allele frequency change in a smaller number of replicates. All significant windows on eigenvectors 1 and 2 are summarised in Tables S2-3.

### Empirical Results: High- (HP) and Low-Predation (LP) Guppies

The motivation for using this dataset was to detect previously identified (Whiting et al., 2021) and novel regions associated with repeated HP-LP adaptation in guppies. AF-vapeR was run over WGS data from five HP-LP population pairs (Figure 5A), each from a separate river, using window sizes of 25, 50, 100, 200 SNPs. Variable window sizes were used to evaluate how robust outliers were to changes in this parameter. Median physical window sizes were 4.8, 10.3, 21.4, and 43.9 kb respectively. We took windows that exceeded null cut-offs for at least two window sizes to increase confidence in candidates, and to ensure spurious signals linked to window size choice were removed. For window sizes of 25 and 50 SNPs we used a quantile cut-off of 99.9%, but relaxed this to 99% for 100 and 200 SNP windows due to the lower overall number of windows and reduced genome-wide variance (Figure 5B-C).

**Figure 5:**
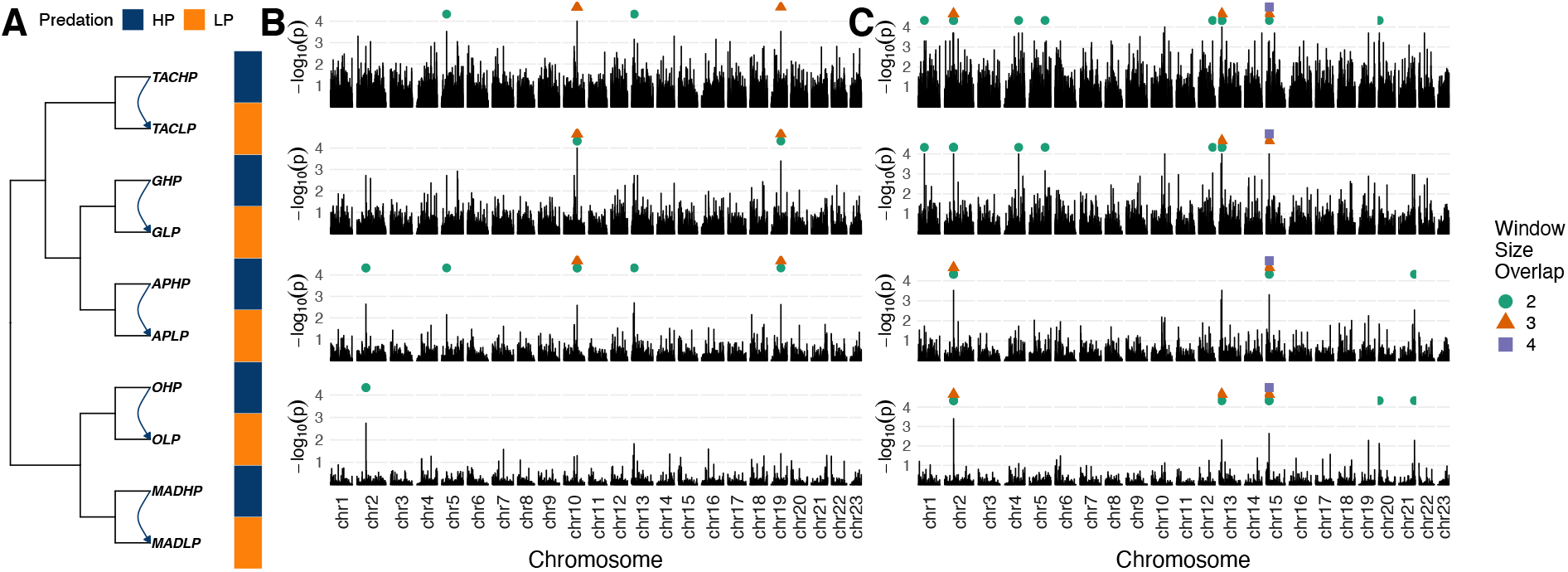
AF-vapeR results for WGS data from HP-LP guppies. Each AFV was calculated between a distinct HP and its corresponding LP population within five rivers in a “replicate pairs” sampling design as illustrated in **A**. Each row in panels **B** and **C** shows genome-wide empirical p-values calculated based on 10,000 null permutations and window sizes of 25 (top), 50, 100 and 200 (bottom) SNPs for primary (**B**) and secondary (**C**) eigenvalues. Agreement among window sizes is shown by points above peaks, with point shape and colour indicative of overlap among window sizes.

This analysis highlighted six windows across five regions of the genome with significant primary eigenvalues on chromosomes 2, 5, 10, 13 and 19 (Table S4). However, all of these regions represented a combination of parallel and antiparallel vectors (Table S4), suggesting that whilst all population pairs exhibit allele frequency differentiation along a common axis, the direction of change was not always consistent among replicates. The regions with the most parallel changes (i.e. only a single anti-parallel replicate to four parallel replicates) were chr5:8127850-8134640 (APHP-APLP antiparallel), and chr10:21021152-21034438 (TACHP-TACLP antiparallel). The results from the primary eigenvalue thus agree with the previous analyses of these populations, in that full-parallelism is limited within this system (Whiting et al., 2021).

On the second eigenvector we identified 13 significant windows over nine regions (Table S5, Figure 5C). Large summed eigenvalues can be caused by either a large eigenvalue 1 plus a small eigenvalue 2, or by a more balanced eigenvalue 1 and 2. The latter are better indicators of multiple parallel axes of allele frequency change. Windows on chr1, chr15, chr20 and chr21 exhibited second eigenvalues accounting for at least 30% of total variance (Table S5). Of these, the window on chr21 was the only window where one population pair exhibited no parallelism with any of the other population pairs. Population loadings onto eigenvectors 1 and 2 (Table S4) highlighted that two of these windows (chr1:9791276-9798829 and chr20:1248797-1296686) show particularly interesting distinct phylogenetic axes of parallelism for the TAC-G-AP and O-MAD lineages (Figure 5A).

The window chr20:1248797-1296686 (Figure S8) intersects with the major candidate previously identified as evolving convergently in the three Caroni rivers (Whiting et al., 2021), and the window chr15:5028361-5066375 (Figure S9) has been demonstrated to evolve in parallel following experimental translocations of HP guppies to LP environments (van der Zee et al., 2021). Neither of these windows exhibited evidence of parallelism from all five rivers, as shown here, and in our previous analysis (Whiting et al., 2021). An in-depth assessment of candidate windows is in the supplementary results.

In conclusion, these results confirm previous analyses that full parallelism is likely limited in this system, whilst also identifying novel candidates and extending previous candidates to more populations that are evolving under a multi-parallel scenario.

### Empirical Results: Galapagos Finch beak morphology

The goal here was to explore an inter-species dataset to examine limitations associated with divergence over time; the common ancestor of the Galapagos finch species group arrived at the islands ~1.5 million years ago (Lamichhaney et al., 2015). The original study used WGS to investigate beak morphology among Galapagos finches (Han et al., 2017) and included 12 between-species comparisons in a multi-comparison sampling design. The focal beak traits of interest were size (small [S] vs large [L]) and shape (pointed [P] vs blunt [B]). These 12 comparisons included all combinations of beak phenotypes, e.g S-S, S-L, L-L, P-P, P-B, B-B (Table 2 in (Han et al., 2017)). We first used all 12 comparisons with AF-vapeR to fully replicate the original study, but expectedly given the range of phenotype comparisons being made, we observed no signals of parallelism. This emphasises the need to carefully consider the input vectors prior to analysis. We next looked at targeted phenotype comparisons, i.e. S-L (six comparisons, nine species) and P-B (five comparisons, seven species) (Figures 6A and 6D). We applied four window sizes (50, 100, 200, 500 SNPs), that were selected due to greater SNP density in this cross-species dataset. Windows reflected median physical sizes of 5.4, 11.1, 22.6 and 57.2 kb. Signatures of parallelism and eigenvalue variance were generally greater in this dataset than in the guppy dataset, so we used stricter outlier criteria to examine only the candidate regions with the strongest evidence (>99.9% of all permuted null values across all window sizes).

**Figure 6:**
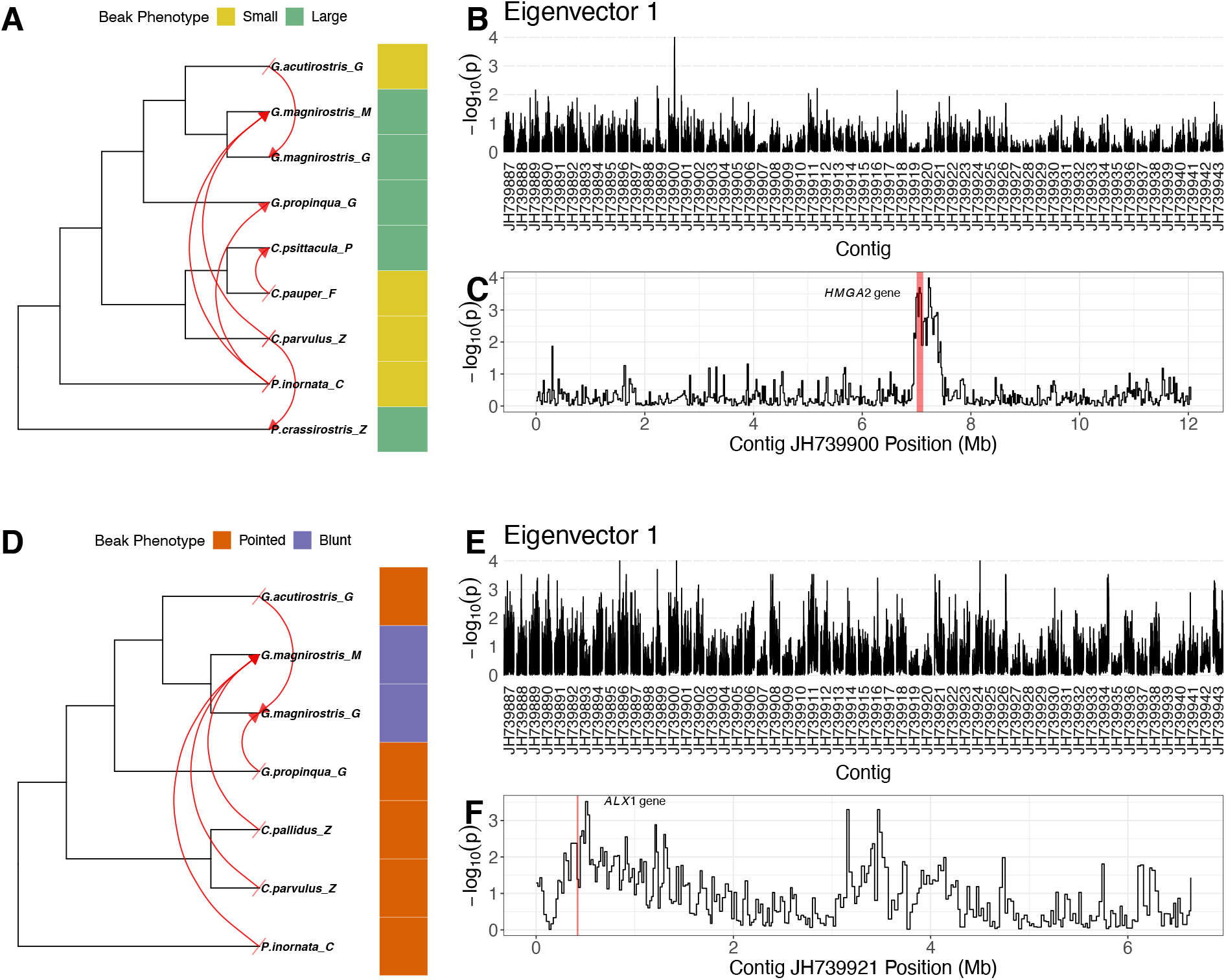
AF-vapeR results for WGS data from Galapagos finches. AFV were calculated between species with contrasting beak phenotypes (small to large or pointed to blunt), with species used multiple times in a multi-comparison design (**A**,**D**). Empirical p-values calculated based on 10,000 null permutations and a window size of 500 SNPs for primary eigenvectors (**B**) highlighting a single strong signature of full-parallelism on contig JH739900 associated with the HMGA2 locus for beak size (**C**). The comparable figure for beak shape (**E**) highlights several loci exhibiting full parallelism, including a window close to, but not including, the candidate ALX1 gene (marked in red) on contig JH739921 (**F**).

A single clear peak of full-parallelism was observed for S-L comparisons, clustered around the *HMGA2* locus and neighbouring genes on contig JH739900 (chromosome 1A) (Figure 6B-C) (Table S6). There were no signals of multi-parallelism on the second eigenvector for S-L vectors. This strong signal of full-parallelism at the *HMGA2* region implies a single axis describing allele frequency change from small to large beak phenotypes; indicative of single S and L haplotype groups. Indeed, clustered genotypes over the two windows overlapping *HMGA2* illustrate cross-species clustering within S and L haplotypes (Figure S10). These results confirm that the evolution of beak size within the galapagos finch radiation has involved the segregation of commonly inherited S and L haplotypes at the *HMGA2* gene among species (Lamichhaney et al., 2016), and no other regions of the genome show this process.

In contrast, for P-B comparisons we observed signals of full-parallelism on fourteen contigs (Table S7-8). The *ALX1* gene has been identified as the primary candidate of interest in the evolution of blunt beak shape (Lamichhaney et al., 2015). Our full-parallelism outliers included a peak adjacent to, but not including, the *ALX1* locus on contig JH739921 (Figure 6E-F), but within the 130kb downstream region identified by Lamichhaney et al. (2015). A lack of parallelism over the *ALX1* gene specifically reflected strong haplotype structure among pointed beak species; genotype clustering suggests four monophyletic P-haplotype groups over the *ALX1* gene (Figure S11). This suggests divergent starting points prevent the identification of parallel P-B axes, and highlight that signatures of full-parallelism erode over time. Inferences may therefore be limited when applying our method to haplotypes with ancient origins.

In total, 16/31 windows (500 SNPs, ± 50kb) with strong signals of full-parallelism contained previously identified and novel candidate genes for beak morphology in this system (discussed fully in supplementary results and Table S9). Of particular note was a signal of full-parallelism detected at *IGF1*, which has been linked to bill morphology in black-bellied seedcrackers (*Pyrenestes ostrinus*) (vonHoldt et al., 2018). In the seedcracker study, no evidence was found to link this gene to beak morphology in *Geospiza*, but our analysis suggests this gene may be important for both taxa.

## DISCUSSION

### Summary of approach

We present AF-vapeR, a novel genome scan method to explore SNP data for signals of parallel evolution based on multivariate allele frequency vectors. Simulating different evolutionary scenarios demonstrates low FPR and considerably reduced FNR relative to common F_ST_ outlier overlap approaches, whilst also distinguishing among different modes of evolution. Empirical analyses demonstrate the ability to identify previously-known candidates of parallelism and to uncover additional candidates. The reduction in Type-II error is derived from re-ordering the workflow to scan for signatures of parallelism explicitly rather than of local adaptation. Identified candidates for parallel evolution can later be assessed for the relative strength of selection or differentiation if desired.

AF-vapeR is analogous to clustering approaches such as cluster separation score (CSS) (as applied by (Jones et al., 2012; Morales et al., 2019; Roberts Kingman et al., 2021)), Cochran–Mantel–Haenszel (CMH) tests (as applied by (Barghi et al., 2019; Orozco-terWengel et al., 2012)), and the C2 statistic (Olazcuaga et al., 2020) (an extension of BayPass’ covariate model to binary groups). Importantly, each of these assumes a single possible axis for parallelism, whereas AF-vapeR can detect multiple axes of parallelism reflecting different trajectories of allele frequency change among replicates. In addition, CSS tests for low within-ecotype/habitat/phenotype distances relative to between, which is true of some cases of full-parallelism, but not all cases as defined here and detected by AF-vapeR. Specifically, AF-vapeR detects similarity in direction of trajectory, as opposed to clustering of starting and endpoints, so both clustered and unclustered cases of full-parallelism (Figure S1) can be detected. Further, our permutation approach to deriving the null distribution incorporates sampling design, and therefore is flexible to sampling designs involving biased group membership (such as a single common ancestor vs many diverged replicates) that may affect other clustering approaches.

AF-vapeR, however, is not an explicit test for selection, but for parallel allele frequency differences. This assumes highly parallel allele frequency change is an indicator of selection through repeated, independent sorting of alleles (but see (Bailey et al., 2015)), but parallelism could arise by other means. For instance, regions linked to blunt beaks in finches parsimoniously evolved once in the common ancestor and were inherited by both monophyletic blunt-beaked species. This sampling design is also likely why baseline P-B eigenvalue 1 scores are high, as there is a genome-wide signal of shared inheritance within the blunt-beaked lineage. To compensate for this, we used stricter cutoffs in the finches compared with stickleback and guppies. Generally, we advise to keep the underlying phylogeny in mind when interpreting results. Hybridisation from an unstudied lineage may also produce parallel allele frequency change, given heterogeneity in the success of introgression across the genome, as suggested for flycatchers (Ellegren et al., 2012) and anchovies (Le Moan et al., 2016). This should not, however, be confused with hybridisation providing adaptive alleles to multiple lineages that are subsequently selected independently. This latter process, observed for example in *Heliconius* (Edelman et al., 2019), is a genuine representation of parallel evolution given increases in the frequency of introgressed variants occur independently (Lee & Coop, 2019).

Recombination rate landscapes commonly obscure genome scans (Lotterhos, 2019), either by inflating variance in divergence, producing longer tails in low-recombination regions that may be falsely identified as signals of selection (Booker et al., 2020), or through inverse associations with background selection (Burri, 2017). Increased variance with reduced recombination is also observed with AF-vapeR (Figure S7). Some resilience to recombination is expected by using SNP count, rather than physical size, windows, given SNP density is linked to recombination landscape (Burri et al., 2015; Stankowski et al., 2019). We therefore expect larger, sparser physical windows in low-recombination regions, and smaller, denser windows in high-recombination regions. This balances the risk of oversampling and having many non-independent windows in tight-linkage in low-recombination regions, and vice-versa. Secondly, because our analysis involves directionality, we can rule out windows with high eigenvalues with parallel and antiparallel vectors. If low recombination inflates variance randomly, it is not expected to produce allele frequency change in the same direction, and additional independent lineages will increase analysis robustness.

Importantly, regions showing strong parallelism may not be under the strongest selection, but may reflect processes governing repeatability specifically, including: heterogeneity in the mutation landscape (Bailey et al., 2018); differential rates of ancient haplotype maintenance (Nelson & Cresko, 2018; Wang et al., 2019); or gene/QTL density landscape (Roberts Kingman et al., 2021). Identifying these regions will help elucidate the influences of these processes on repeatability.

### Recommendations

We recommend a minimum of four replicate vectors to explore full- and multi-parallelism. Because maximum eigenvalues are constrained to *m* (for *m* AFV), increasing *m* increases the distinctiveness of eigenvalue peaks and reduces the likelihood of false-positives. Beyond studies of repeatability, this approach may be well-suited to evolve-and-resequence experiments (e.g. (Barghi et al., 2019; Rudman et al., 2021; Tusso et al., 2021)), where replication is common and allele frequency change may be moderate and recorded after only a few generations. Because patterns of parallelism break down over time, seen here with *ALX1* haplotypes in finches, our approach is best-suited to intra-specific comparisons, or those among relatively young clades. Whole-genome data is desirable, but not a prerequisite (e.g. RAD-seq shown here), and given allele frequencies are the focus pool-sequencing data could also be used. Whilst the permutation approach accounts for sampling design, some sampling designs may bias eigenvalues more than others, particularly repeatedly sampling the same population when calculating AFV. In such cases, greater stringency can be applied when calling outliers, as demonstrated here when analysing the guppy dataset compared with the finch dataset. Datasets used here were filtered with a minor allele frequency (MAF) filter of 5%, but 1% may be sufficient. MAF-filtering is important so that SNP windows are not overpopulated by many SNPs carrying minimal information.

Visualising trees and genotypes at candidate regions can be informative in understanding how replicates load onto eigenvectors and how populations cluster within parallel or multi-parallel clusters (e.g. Figure S8-11). Further, per-SNP scores available in each window’s projected **A** matrix can be used to identify SNPs whose allele frequencies are driving variance (e.g. Figure S8). Examining individual SNP scores is akin to examining the loading of variables during principal component analysis (PCA) to assess which variables are best reflected in each principal component. We reiterate that AF-vapeR cannot detect cases where all replicates divergently modify the same gene or locus (molecular convergence; Figure 2C), as this produces geometric non-parallelism. Such regions may still be of interest, however, and combining AF-vapeR with selection scans may highlight such regions.

The study of repeatable evolution in the genome is increasingly popular in natural systems and in experimental evolution. Progression of this field is essential for better understanding the determinism, predictability, and contingencies linked with adaptation, with implications for issues including rapid adaptation to changing environments (Reid et al., 2016) and the evolution of antibiotic resistance (A. C. Palmer & Kishony, 2013). There are also other biological questions where the goal is similarly to identify shared haplotypes among groups, such as sex-linked loci in the heterogametic sex (D. H. Palmer et al., 2019). Despite this, there remain limited approaches tailored towards detecting signatures of repeatability from genomic data. Our method requires only a standard VCF file, a population map, and a description of vectors to calculate, and should be accessible to researchers familiar with common multivariate analyses such as PCA or principal coordinate analysis (PCoA). We therefore expect AF-vapeR to become an important tool for detecting convergence alongside currently available approaches.

## ACKNOWLEDGEMENTS

The authors would like to thank Matthew Webster (Uppsala University), and Andrew MacColl (University of Nottingham) and Isabel Santos Magalhaes (University of Roehampton) for insightful discussions regarding the finch and stickleback datasets respectively. The authors also thank members of the University of Exeter’s BS-popgen group and Sam Yeaman (University of Calgary) and members of the Yeaman lab for providing feedback on the manuscript and project. HPC infrastructure support was provided by The University of Exeter’s High-Performance Computing (HPC) facility (ISCA).

## DATA AVAILABILITY

AF-vapeR has been made available for download at github (https://github.com/JimWhiting91/afvaper). The code available for simulations and analyses of simulations and empirical datasets is available at (https://github.com/JimWhiting91/afvaper_analyses_ms). Sequencing reads associated with the stickleback datasets are available at the European Nucleotide Archive (ENA) (accession: PRJEB20851). Sequencing reads associated with the guppy dataset are available at the ENA (accessions: PRJEB43917 and PRJEB10680) and the VCF used is available from FigShare (doi:10.6084/m9.figshare.14315771). The VCF for the galapagos finch data was acquired from Dryad (doi: 10.5061/dryad.7155d).

## AUTHOR CONTRIBUTIONS

JRW and MJvZ conceived the project. JRW developed the software, and performed simulations, analyses and figure production. JRP assisted software development through testing and simulation development. JRW, JRP and BAF were involved in writing the manuscript. BAF supervised the project.

## CONFLICTS OF INTEREST

The authors declare no conflicts of interest.

## Notes

https://github.com/JimWhiting91/afvaper

https://github.com/JimWhiting91/afvaper_analyses_ms

### Competing Interest Statement

The authors have declared no competing interest.

### Summary of Updates

Manuscript updated to be shortened and improve readability. Some information has been added to the supplementary materials, including new supplementary figures. No new results or changes to the content have been made.

