## Supplementary material for "AF-vapeR: A multivariate genome scan for detecting parallel evolution using allele frequency change vectors": Figure S

### Supporting Material - Supplementary Methods

#### Full simulation methods

For all simulations, we assembled a metapopulation with an ancestral founding population and four daughter populations (Figure S2). Each daughter population exchanges migrants with the ancestral founding population, such that each daughter population receives a proportion of M individuals each generation from the founding population, and the founding population receives a proportion of M/4 individuals from each of the four daughters (for a total proportion of M new migrants). We then expose the metapopulation to three different scenarios in which evolution occurred at a focal mutation: 1) 'Full Parallel' - All daughters receive the same mutation during their founding and experience positive selection on this common variant; 2) 'Multi Parallel' - Two mutations arise in the ancestral population, one of which is inherited by one pair of daughters and the other is inherited by the remaining pair, and all four daughters experience selection on their respective mutation; 3) Divergent: Four mutations arise in the founding population and each daughter receives and experiences selection on its own mutation. In all cases, selection takes place with a selection coefficient (s) of either 0.01, 0.05 or 0.1 that is positive in daughters and negative in the founding population to promote divergence. Migration (M) occurs at a rate of 0, 0.0025 or 0.01 between the founding population and daughters. Finally, the population size of daughters (DP) was manipulated to vary between 400, 2,500 and 10,000 individuals; founding populations always comprised 10,000 individuals. These three varying parameters (s, DP, M) were combined factorially to yield 27 parameter sets, each of which was simulated with 100 iterations, and across all three evolutionary scenarios. Following the founding of daughter populations, simulations proceeded for 200 generations to allow for divergence to occur. In the event of mutations being lost in any of the four daughter populations, simulations were returned to the point at which daughters had been founded and restarted with a new seed. All parameters are summarised in Table S1.

These 27 scenarios were selected to explore where popular genome scans and summary statistics are predicted to struggle in detecting genetic parallelism/convergence, particularly with respect to migration-selection balance. Under low DP values, we expect daughters to experience more drift at neutral loci which may obscure scans based on differentiation outliers. This effect is exacerbated at low or zero M where drifting loci cannot be shared between daughters and their founding population. At high rates of M, adaptation may be constrained to only weak or subtle shifts in allele frequencies, an effect which is strongest when selection is weak. Indeed, the choice of a maximum M of 0.01 and minimum s of 0.01 is to explicitly simulate the point at which migration-selection balance is equal and adaptation should be severely limited.

Simulations took place on a 20 Mb chromosome, with the focal site at 10.0 Mb, and a constant recombination rate of 1e-8. We also explored the effects of variable recombination, in which recombination increased 100-fold linearly from a low of 1e-9 to a high of 1e-7 along a 22 Mb chromosome, with recombination increases every 2 Mb. With variable recombination, the focal site for selected variants occurred at 1 Mb (r = 1e-9), 11 Mb (r = 1e-8) and 21 Mb (r = 1e-7), and we ran 25 iterations at fixed parameters of s = 0.05, m = 0.0025 and DP = 10,000 at low, medium, and high recombination rates (total of 75 iterations).

Following an additional 200 generations of selection, tree sequences (Haller et al., 2019) were exported from SLiM, recapitated, and converted to VCF files of 20 randomly selected individuals per population (total of 100 per VCF) using pyslim. For each VCF, we then calculated FST between each of the daughter populations and the founding population in 10kb windows with vcftools (v0.1.16 (Danecek et al., 2011)). We then used AF-vapeR with window sizes of 50, 200, and 500 SNPs to scan each iteration for signals of parallel evolution at the focal site. For each parameter set, we calculated the false positive (FPR) and false negative rate (FNR). For each iteration, we calculated the FPR as the number of detected signals of parallel evolution by AF-vapeR (compared against null quantiles of α=0.95, α=0.99 and α=0.999), or overlapping FST outlier windows (overlapping in all four, any three, or any two, outlier sets above the same quantile cut-offs) at neutral loci (beyond an exclusion zone of 0.1 cM around the focal site). We then averaged FPR across all 100 iterations to estimate FPR of each parameter set. For FNR, we counted the number of iterations in which parallel evolution or overlapping outlier FST windows were not detected within a region of 0.01 cM around the focal site at each null quantile, and divided this by the number of iterations (100).

#### Supporting Material -Supplementary Results

##### Stickleback candidates

The following windows were identified as fully parallel above the 99.9% cutoff: chromosome 1 (21721603-21849857) included the *atp1a1* gene known to be important for freshwater adaptation (Nelson & Cresko, 2018); chromosome 5 (4273793-4332432) is between several QTL associated with marine-freshwater feeding (Erickson et al., 2016; Miller et al., 2014); chromosome 7 (19280164-19396323) harboured SNPs covering the claudin genes *cldnc* and *cldnh*, the latter of which is differentially expressed between marine and freshwater stickleback exposed to salinity gradients (Gibbons et al., 2017); chromosome 11 (5577181-5794574) included feeding (Glazer et al., 2015) and sensory (Wark et al., 2012) QTL and has been detected in other parallel evolution studies in British Columbia (Hohenlohe et al., 2010); and chromosome 20 (6190327-6247151) includes the genes *ddc* and *grb10b*, the former of which is suspected to be involved in parallel evolution through copy number evolution (Hirase et al., 2014).

#### Guppy candidates

We will first discuss the candidate window chr20:1248797-1296686. Plotting the allele frequencies of this region highlights the clear Caroni drainage (Tacarigua [TAC]-Guanpo [G]-Aripo [AP]) vs non-Caroni (Oropouche [O]-Madamas [MAD]) haplotype structure (Figure S8B), and also reveals AF change in the same direction for HP > LP within these two groups. Examining the scores of individual SNPs within this window shows that both eigenvectors share a common high-loading SNP (denoted by an arrow in Figure S8). This SNP intersects the only predicted gene in this region (ENSPREG00000013524), and indicates that both haplotypes may have independently acquired the same mutation, which is under parallel selection in all rivers. Interestingly, using a protein sequence blast against NCBI's nr database we find that this gene exhibits sequence similarity with various teleost syncytin genes. This family of proteins are essential for placental development in mammals (Dupressoir et al., 2011), and are expressed in the maternal follicle transcriptome of other live-bearing poeciliids (Guernsey et al., 2020).

Another region of interest is chr15:5028361-5066375, which has been associated with convergent HP-LP evolution elsewhere (Fraser et al., 2015; van der Zee et al., 2021; Whiting et al., 2021). This region of the genome was, uniquely, detected across all four tested window sizes. Again we observe a signature of multi-parallelism here in which all five rivers loaded onto one, or both eigenvectors, but with no clear ancestral structure (Table S4). Rather than reflecting the underlying phylogeny, this is suggestive of complex haplotype structure, such that there may be two, or more, HP/LP-adapted haplotypes segregating in the same direction between HP and LP populations in all lineages. Clustering of haplotypes (Figure S9) within this window highlights that eigenvector 1 represents a change in the frequency of an HP-associated haplotype found in Madamas, Oropouche and Tacarigua, whilst eigenvector 2 reflects the change in the frequency of an LP-associated haplotype found in all rivers except for Oropouche, which carries a novel LP haplotype associated with a deletion (Whiting et al., 2021). Both eigenvectors had high-loading SNPs clustered around the 6th and 7th exons of the *B-cadherin* gene, suggesting this may be a common target for selection through non-parallel mutation.

The window at chr1:9791276-9798829 also exhibited individual SNPs that were fully parallel in all rivers, but clear ancestral haplotype structure associated with Caroni and non-Caroni lineages. The window is intergenic, but ~5kb upstream of *kpna7*, an ovarian-expressed gene important for early embryonic development in mammals (X. Wang et al., 2012) and fish (L. Wang et al., 2014). Given differences in reproductive biology are key repeatable phenotypes among HP and LP guppies, this window and the window on chromosome 20 represent significant candidates of interest regarding these phenotypes.

##### Finch candidates

Beyond the ALX1 locus, we recovered signals of full parallelism at genes previously identified in genome scans of Galapagos finch beaks (STK3, VPS13B, NT5C1B, RDH14, ADK) (Chaves et al., 2016; Lamichhaney et al., 2015; Lawson & Petren, 2017). We also identified windows with novel candidate genes with probable roles in craniofacial development (LRIG3, IGF1, PMCH, PARPBP, NUP37, GDF6, PTDSS1, OSR2, SAMD12, COLEC10, ENPP2, AHSA1, AP3M1, VCL, WNT6, WNT10A, FEV, TP63) (Table S7).

Drainage-structuring of ancestral variation and a common functional pathway shape limited genomic convergence in natural high- and low-predation guppies. *PLoS Genetics*, *17*(5), e1009566.

#### Supporting Material - Supplementary Figures:

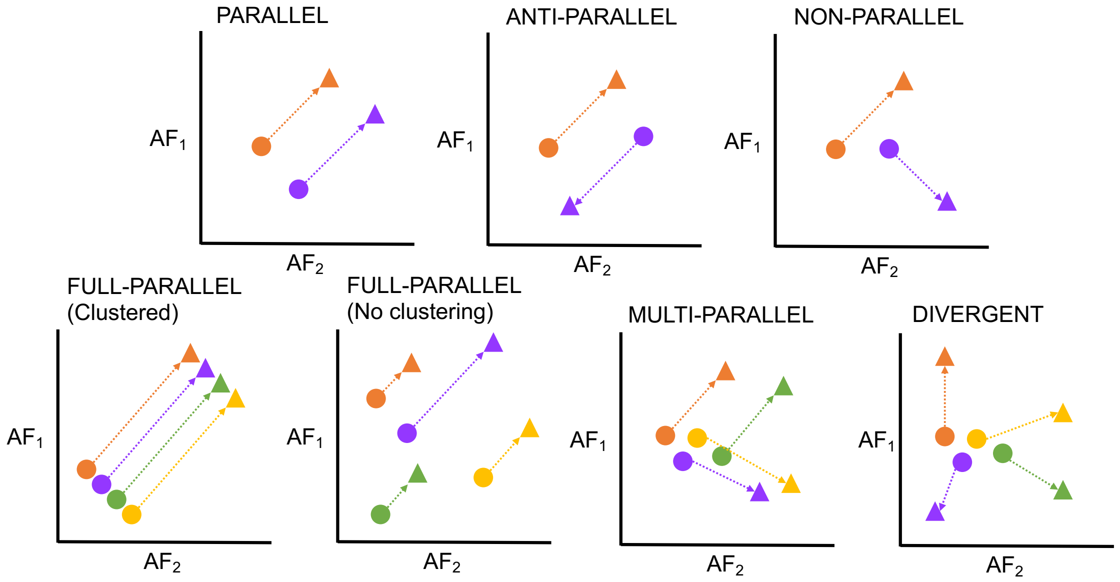


**Figure S1:** Schematic representations of parallelism definitions as recognised by AF-vapeR. All plots are presented in two-dimensional allele frequency (AF) space, and show the change in allele frequencies between a startpoint population (circles) and an endpoint population (triangles). Different colours represent replicate sampling pairs. The first row describes broad terms: "Parallel" (change in the same direction along the same trajectory); "Anti-parallel" (change in the opposite direction along the same trajectory); and "Non-parallel" (change along a different trajectory). The second row visualises different forms of parallelism detected by AF-vapeR among four replicate pairs: "Full-parallel" (all replicates show parallel allele frequency change along the same trajectory), which can be detected regardless of whether start/endpoints are clustered; "Multi-parallel" (two "parallel" trajectories exist, with two replicate population pairs diverging in parallel along each trajectory); "Divergent" (all replicate pairs diverge along non-parallel trajectories).

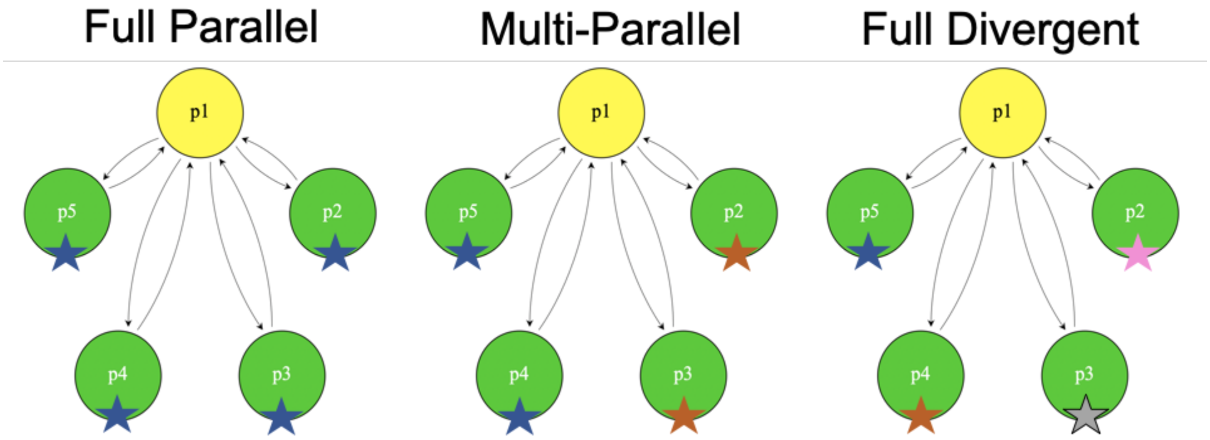


**Figure S2:** Summary of simulation models used to evaluate AF-vapeR under different evolutionary scenarios. All simulations involved a founding population (p1) founding four daughter populations (p2-5), each of which inherits a beneficial mutation (stars) where mutation colour signifies whether populations received the same or different mutations present as standing genetic variation at time of founding. Arrows between p1 and p2-5 signify ongoing reciprocal migration between p1 and its daughters at a rate of *M*, but migration into p1 is *M*/4, such that all populations receive the same proportion of immigrants.

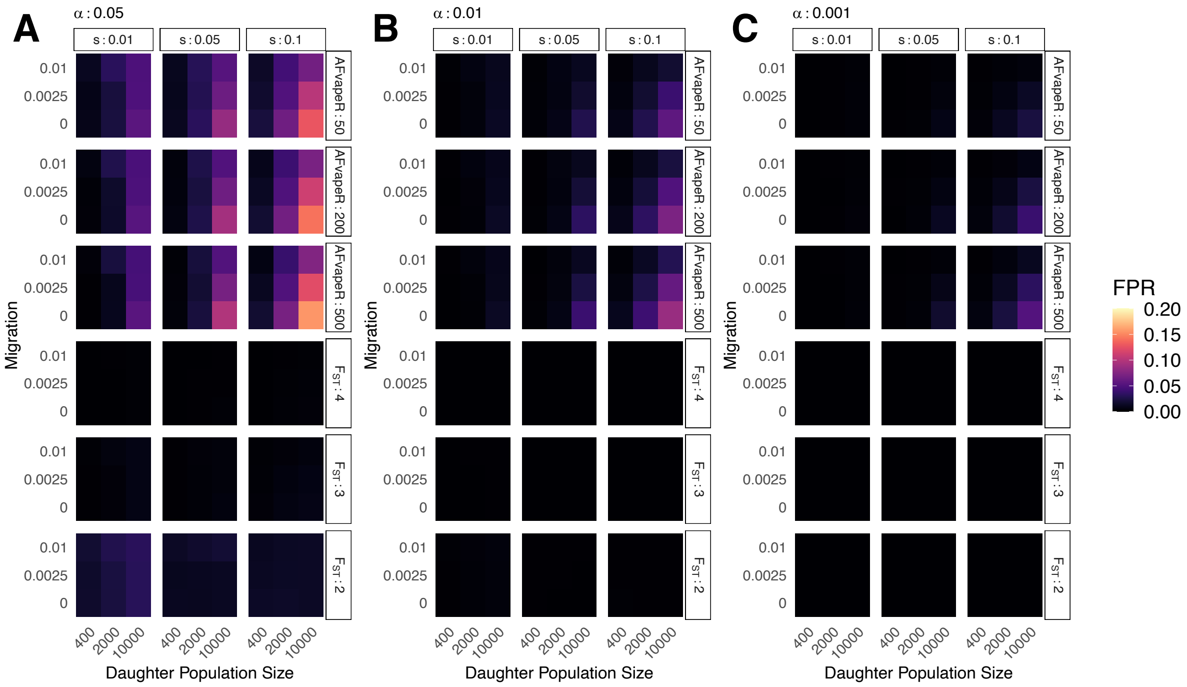


**Figure S3:** False positive rates (FPR) associated with AF-vapeR and comparing $F_{ST}$ outliers tests under a "full-parallelism" simulation. AF-vapeR facet labels correspond to the number of SNPs per window used in the analysis (50, 200, and 500 SNPs). $F_{ST}$ facet labels correspond to the number of overlapping populations required to call a region as overlapping, (i.e. 2 = an outlier in any 2 of the 4 populations, 3 = an outlier in any 3, 4 = outlier detected in all four). Panels A, B and C show FNR calculated based on taking outliers at quantile cutoffs of 95%, 99%, and 99.9% respectively. Within facet rows, each grid space represents the FNR averaged over 100 iterations for one of the 27 factorial parameter combinations, where facet columns = strength of selection on beneficial mutation, y = migration rate, and x = daughter population size.

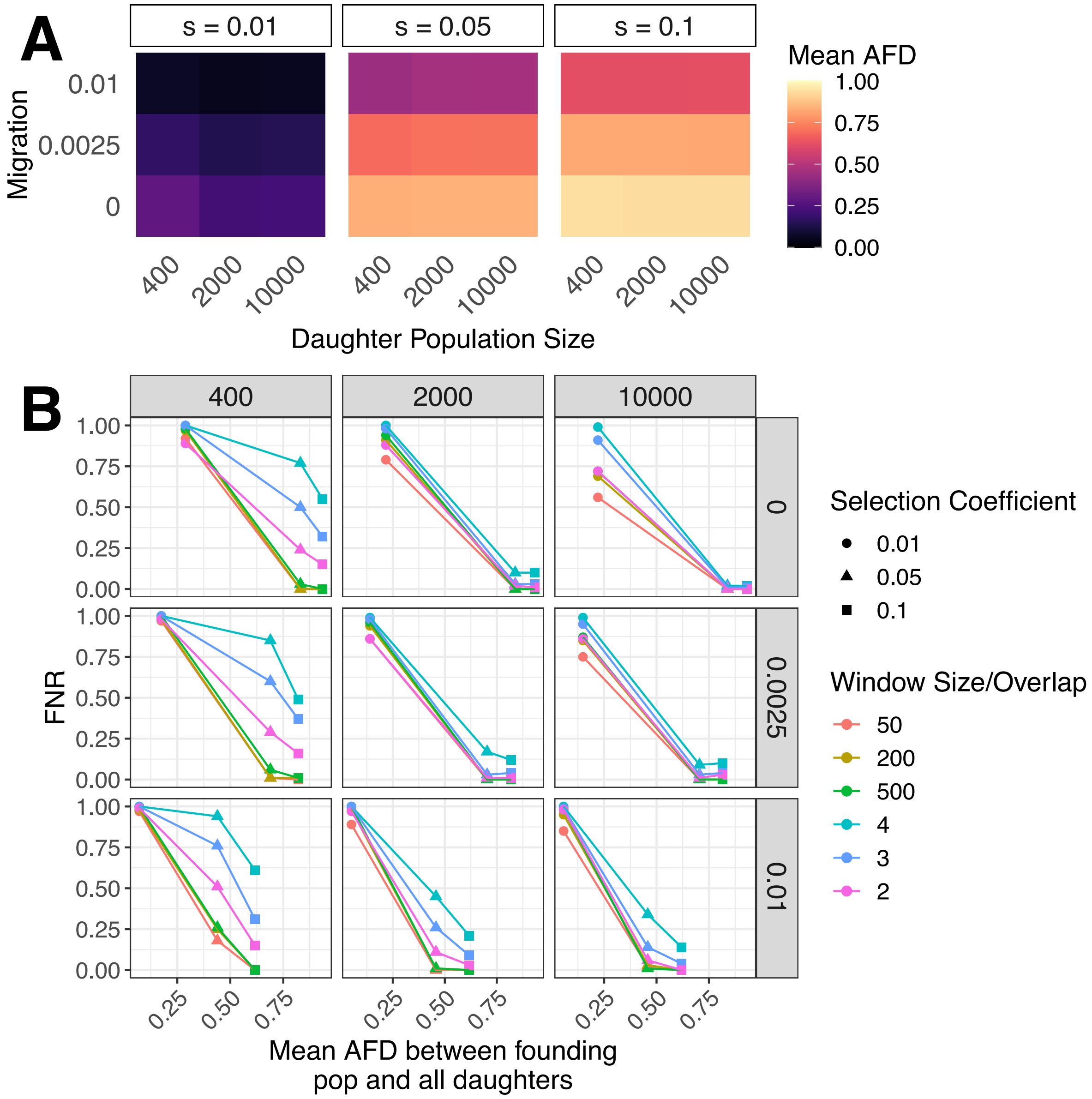


**Figure S4:** The interaction between FNR and allele frequency differentiation (AFD) at the selected site across parameter space. Both migration rate (grid rows) and selection coefficient (facet columns) were the drivers of AFD across the 27 parameter combinations (A). The relationship between FNR and AFD is shown in (B). Facet columns show daughter population size parameter and facet rows show migration rate parameter value. Selection coefficient parameters are denoted as point shape, and point/line colour denotes analysis (AF-vapeR window size = "50", "200", "500"; $F_{ST}$ overlap among "2", "3", or "4" daughter populations). AF-vapeR is more sensitive to AFD, shown as generally steeper reductions in FNR with increasing AFD.

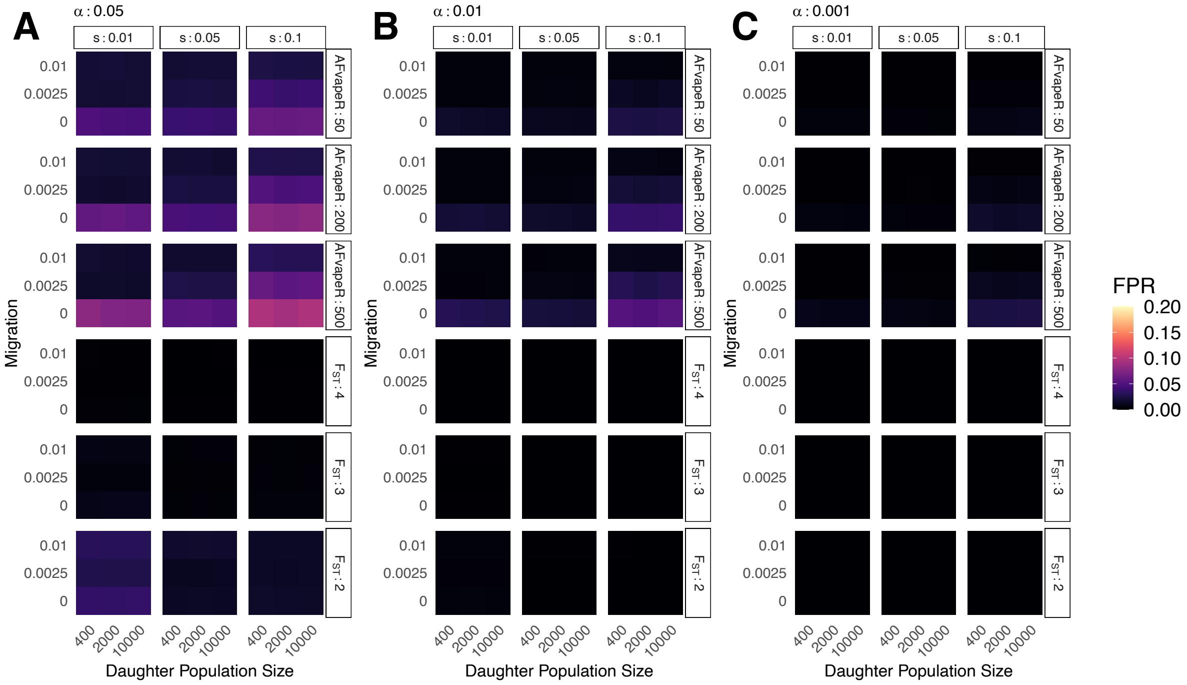


**Figure S5:** False positive rates (FPR) associated with AF-vapeR and comparing $F_{ST}$ outliers tests under a "multi-parallelism" simulation. AF-vapeR facet labels correspond to the number of SNPs per window used in the analysis (50, 200, and 500 SNPs). $F_{ST}$ facet labels correspond to the number of overlapping populations required to call a region as overlapping, (i.e. 2 = an outlier in any 2 of the 4 populations, 3 = an outlier in any 3, 4 = outlier detected in all four). Panels A, B and C show FNR calculated based on taking outliers at quantile cutoffs of 95%, 99%, and 99.9% respectively. Within facet rows, each grid space represents the FNR averaged over 100 iterations for one of the 27 factorial parameter combinations, where facet columns = strength of selection on beneficial mutation, y = migration rate, and x = daughter population size. AF-vapeR outliers are determined on the basis of being detected as outliers on the second eigenvector.

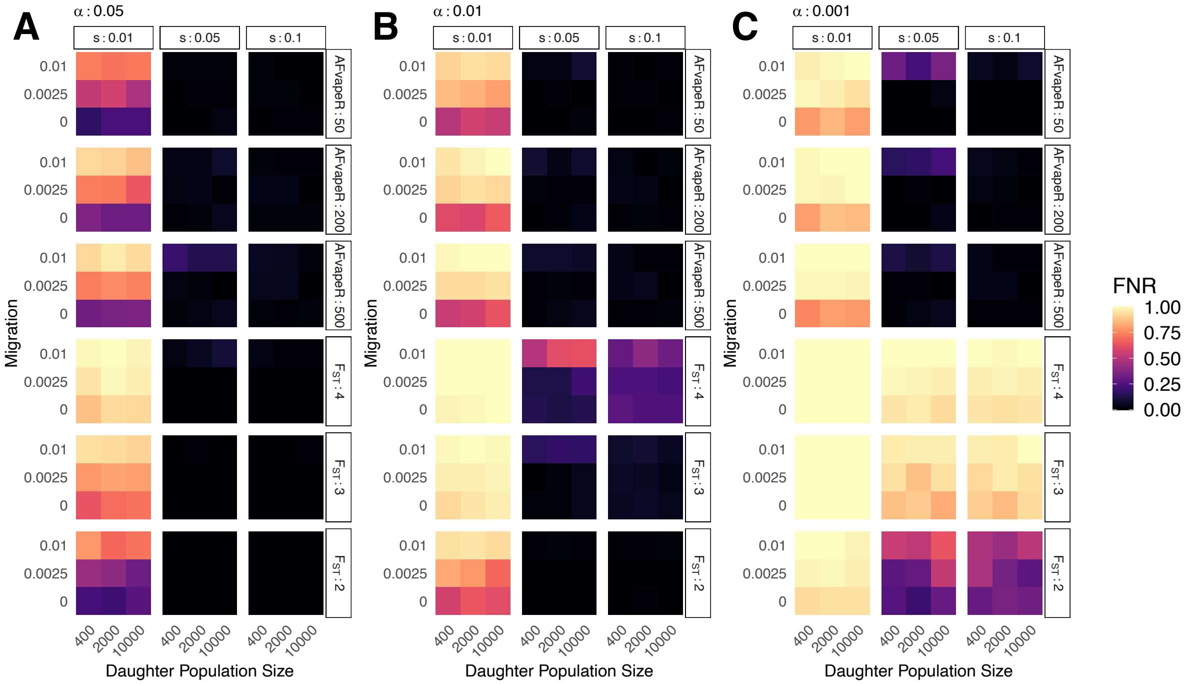


**Figure S6:** False negative rates (FNR) associated with AF-vapeR and comparing $F_{ST}$ outliers tests under a "multi-parallelism" simulation. AF-vapeR facet labels correspond to the number of SNPs per window used in the analysis (50, 200, and 500 SNPs). $F_{ST}$ facet labels correspond to the number of overlapping populations required to call a region as overlapping, (i.e. 2 = an outlier in any 2 of the 4 populations, 3 = an outlier in any 3, 4 = outlier detected in all four). Panels A, B and C show FNR calculated based on taking outliers at quantile cutoffs of 95%, 99%, and 99.9% respectively. Within facet rows, each grid space represents the FNR averaged over 100 iterations for one of the 27 factorial parameter combinations, where facet columns = strength of selection on beneficial mutation, y = migration rate, and x = daughter population size. AF-vapeR outliers are determined on the basis of being detected as outliers on the second eigenvector.

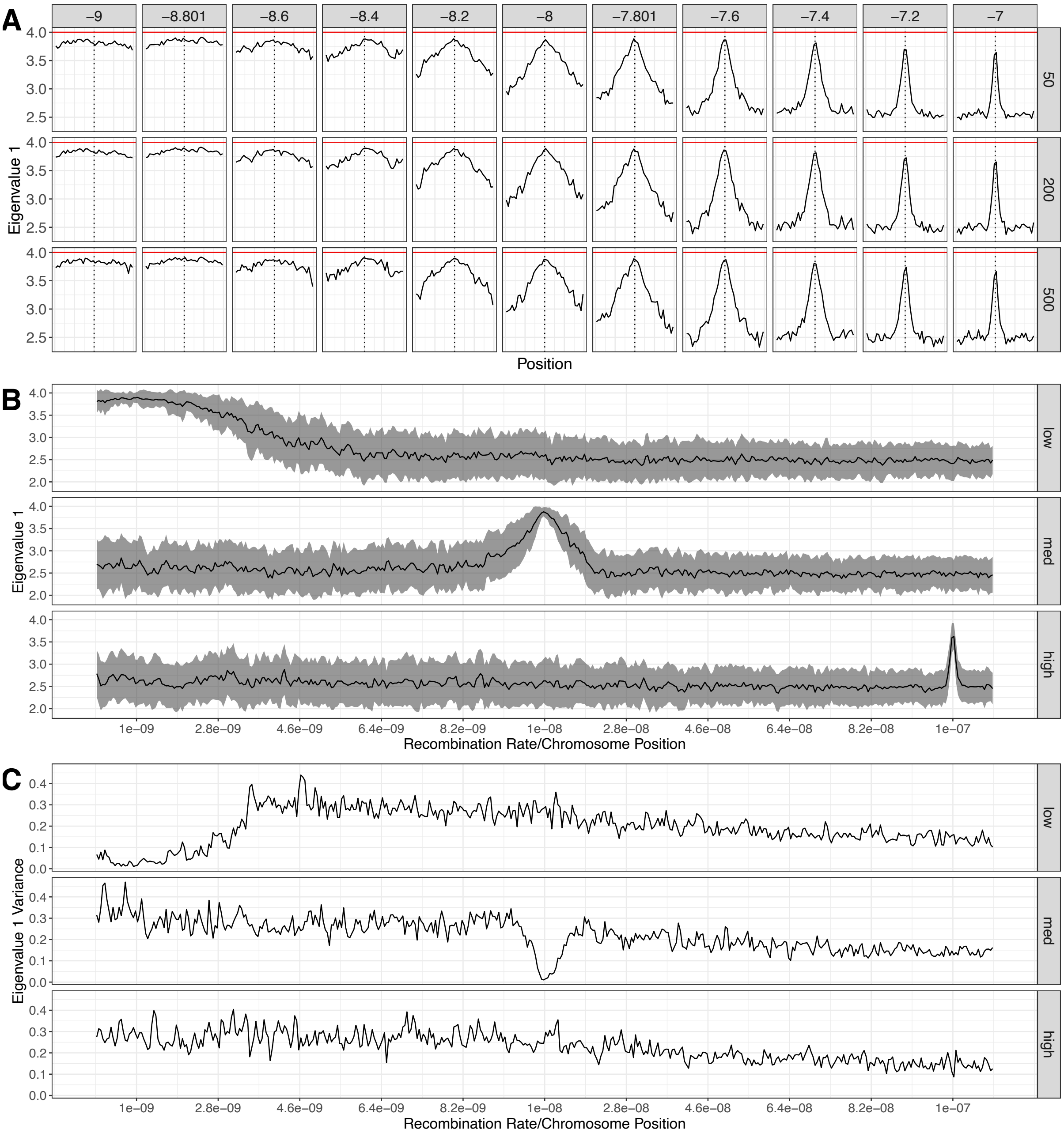


**Figure S7:** Effect of recombination on eigenvalue 1 under full parallelism. (A) The shape of the eigenvalue 1 peak signature around the site under selection is shown across different recombination rates (columns) and different SNP window sizes (rows). The red line denotes the maximum value of 4 (simulations include four daughter populations), and the vertical dashed lines denote the site under selection. (B) Eigenvalue 1 peaks (± sd) along a chromosome with 100-fold variation in recombination. Selected sites were introduced either in a region of low recombination (1e-9), medium recombination (1e-8), or high recombination (1e-7), plotted separately as rows. (C) Eigenvalue 1 variance at neutral is negatively associated with recombination rate across a variable recombination landscape, with recombination increasing from left to right.

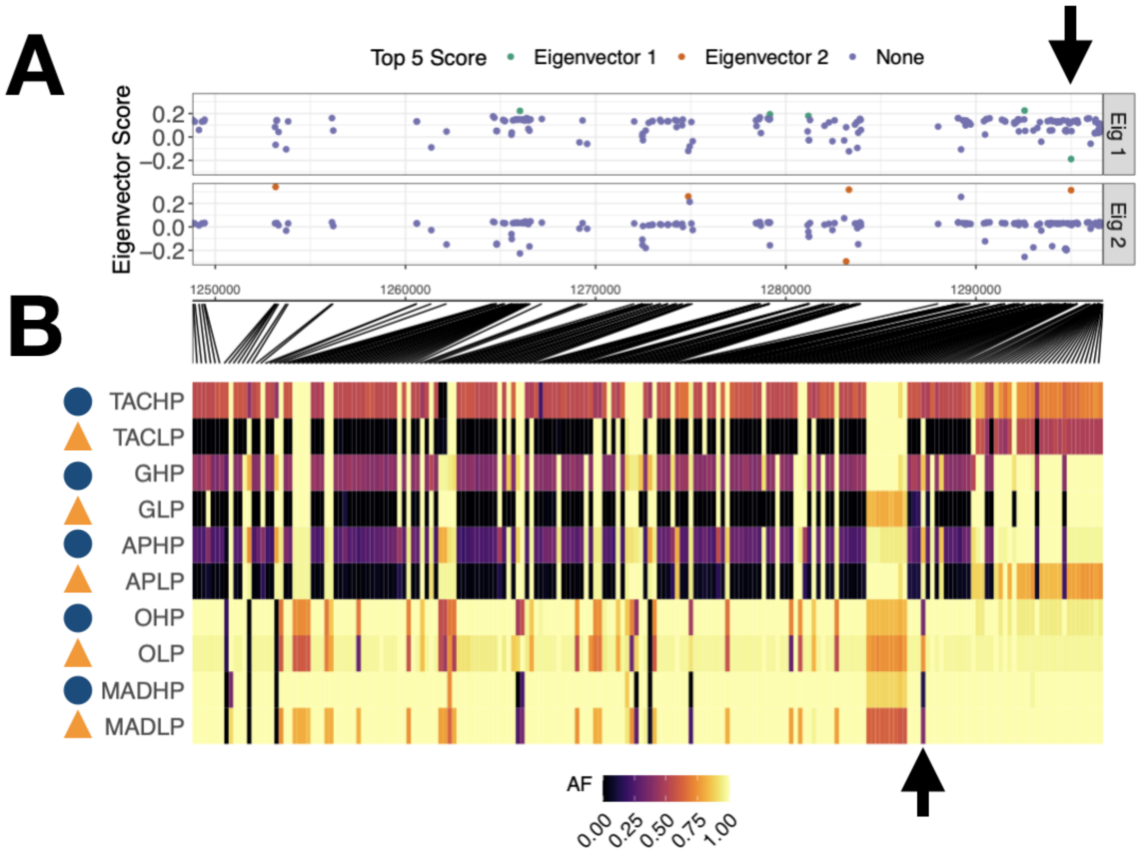


**Figure S8:** Individual SNP scores associated with multi-parallelism within the region chr20:1248797-1296686. In panel (**A**) the SNPs with the top 5 scores on eigenvector 1 and 2 are highlighted in green (Eig1) and orange (Eig2). The raw allele frequencies for all 10 populations are plotted in panel (**B**), with black lines connecting the BP position of each SNP in (**A**) to its position in the allele frequency grid in (**B**). HP and LP populations are paired in rows, and prefixes (TAC, G, AP, O and MAD) denote rivers. The black arrows highlight a single SNP that is in the top 5 scores for both eigenvectors (**A**), is shared in both haplotypes, and increases in frequency in all LP populations (**B**).

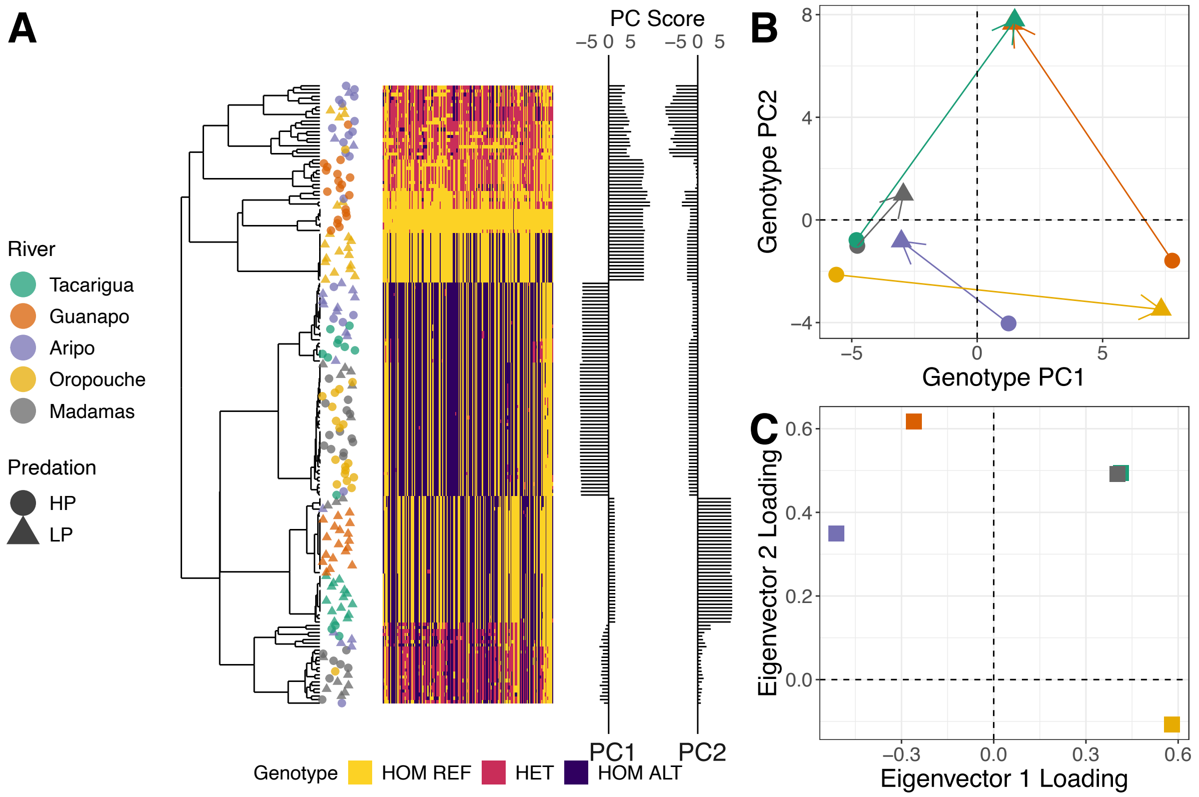


**Figure S9:** An example of multi-parallelism in guppies on chr15. Clustered genotypes (**A**) of all individuals based on the 200 SNPs in the window chr15:5028361-5066375, reveals clustering of LP-associated haplotypes towards the bottom of the dendrogram (high PC2), and a cluster of HP-associated haplotypes in the centre (low PC1). Each row represents the genotypes of an individual, ordered according to the dendrogram derived from clustering of genotypes based on principal component analysis (PCA). Individual PC1 and PC2 scores are shown in-line with plotted genotypes. (**B**) Trajectories of change within genotype PC space within rivers, shown are population centroids. A reduction in allele frequencies of haplotypes in the central cluster 'HP-associated haplotype' can be seen as positive trajectories along Genotype PC1 in three rivers. Positive changes along Genotype PC2 are associated with increased frequency of sites associated with the 'LP-associated haplotypes', seen in four rivers. (**C**) Parallelism among each of these allele frequency change trajectories as captured by the AF-vapeR eigenvectors. Positive eigenvector 1 and 2 loadings reflect the direction and strength of change seen between population-specific centroids in (**B**). Predation phenotype and river are shown as shape and colour of points at dendrogram tips and in (**A**) and (**B**).

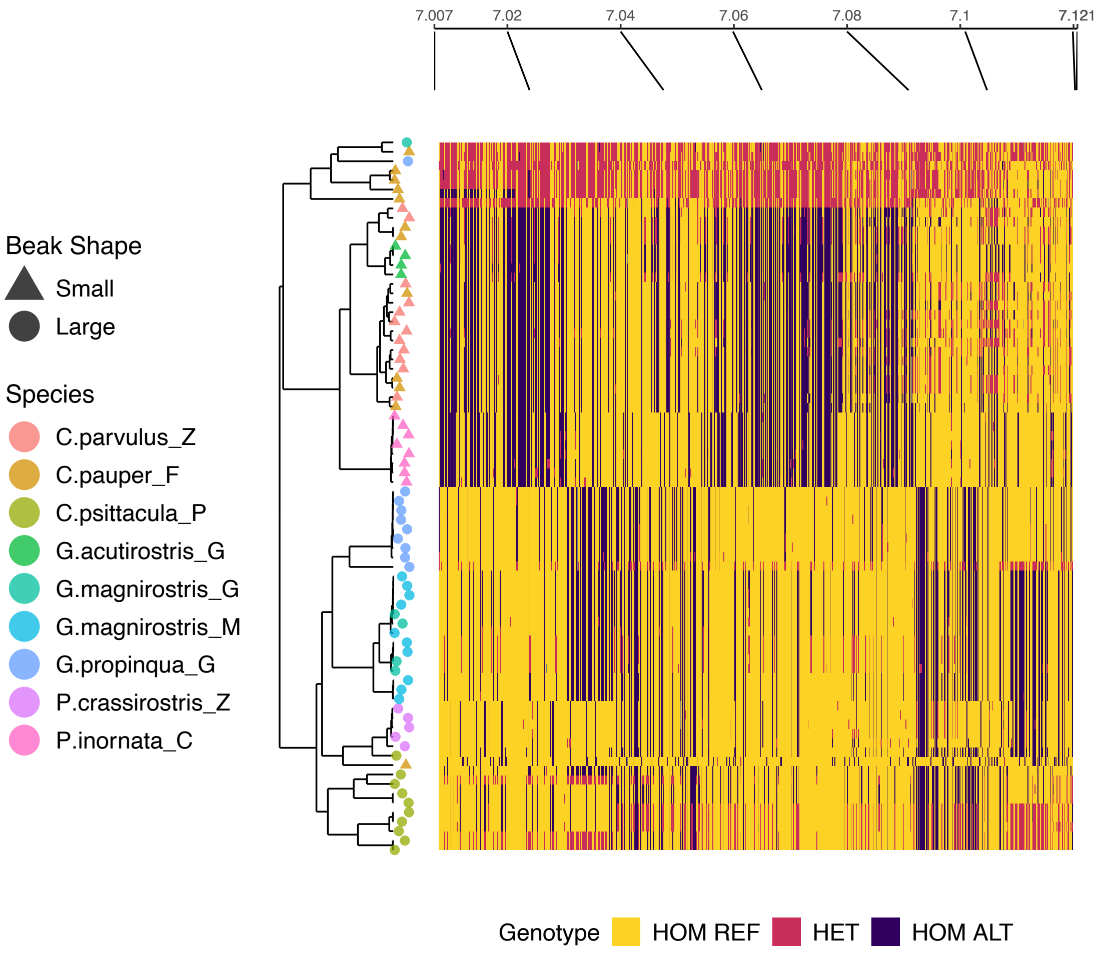


**Figure S10:** Clustered genotype plot within the window including the *HMGA2* gene. Each row corresponds to the genotypes for every SNP in the window JH739900:7007107-7120688 for a given individual. Individuals are clustered according to their haplotype on the y-axis, with the clustering dendrogram shown. At dendrogram tips, species are denoted by point colour, and beak phenotype denoted by point shape. The approximate positions of SNPs every 20kb along the contig are shown at the top.

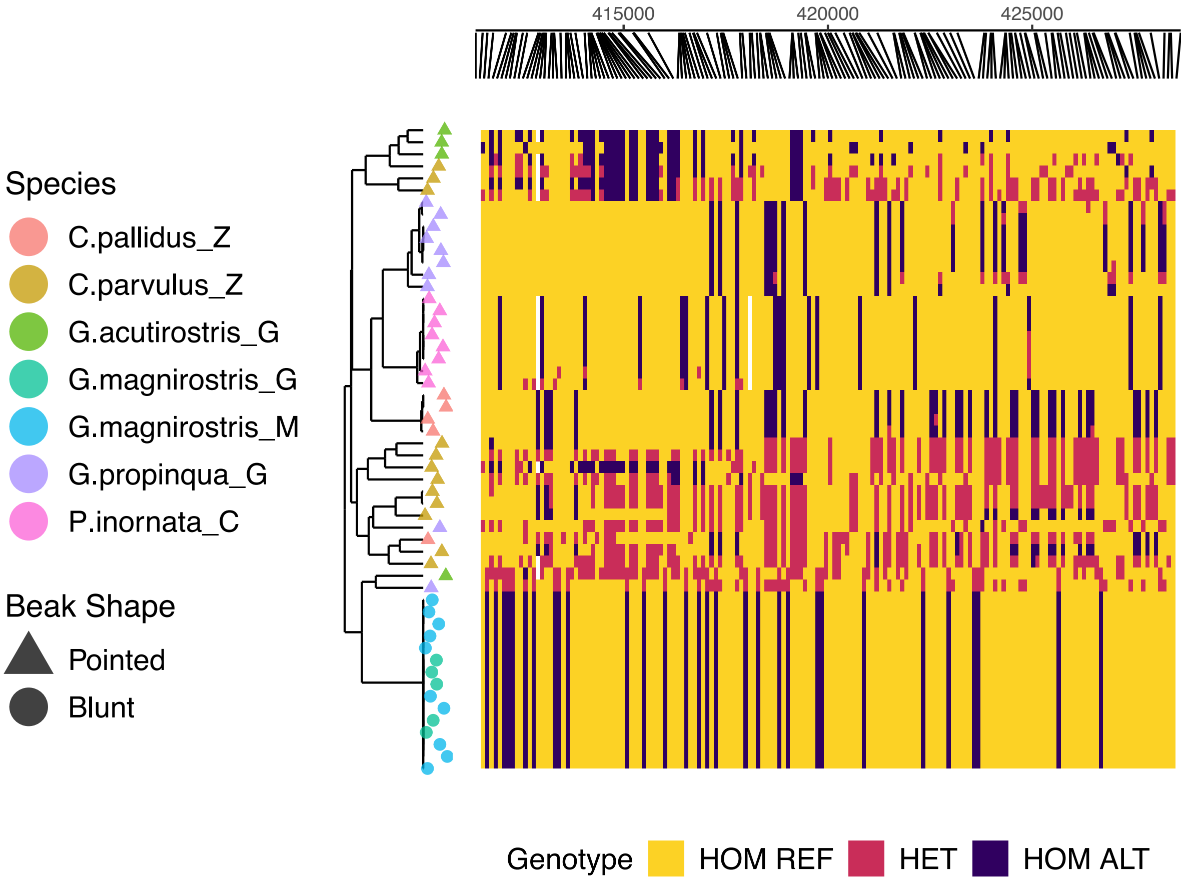


**Figure S11:** Clustered genotype plot within the window including the *ALX1* gene. Each row corresponds to the genotypes for every SNP in the window JH739921:411351-428776 for a given individual. Individuals are clustered according to their haplotype on the y-axis, with the clustering dendrogram shown. At dendrogram tips, species are denoted by point colour, and beak phenotype denoted by point shape. The corresponding positions of every SNP along the contig are shown at the top of the figure.
